# RESTRICT-seq enables time-gated CRISPR screens and uncovers novel epigenetic dependencies of SCC resistance

**DOI:** 10.1101/2025.09.17.676440

**Authors:** Dreyton G Amador, Justin Anthony Powers, Ashley G. Njiru, Zahra Ansari, Yvon Woappi

## Abstract

Cancer cell evasion of therapy is a highly adaptive process that undermines the efficacy of many treatment strategies. A significant milestone in the study of these mechanisms has been the advent of pooled CRISPR knockout screens, which enable high-throughput, genome-wide interrogations of tumor dependencies and synthetic lethal interactions, advancing our understanding of how cancer cells adapt to and evade therapies. However, the utility of this approach diminishes when applied to dynamic biological contexts, where processes are transient and sensitive to routine cell culture manipulations that introduce noise and limit meaningful discoveries. To overcome these limitations, we present RESTRICT-seq, a next-generation pooled screening methodology that restricts Cas9 nuclear activation in controlled, repeated cycles. By confining Cas9 catalytic activity to strict temporal windows, RESTRICT-seq mitigates undesired fitness penalties that accumulate throughout CRISPR screens. When benchmarked against conventional pooled screens and standard inducible CRISPR protocols, RESTRICT-seq revealed significantly fewer divergent cell clones and increased signal-to-noise ratio, overcoming a key limitation of traditional methods. Leveraging RESTRICT-seq, we conducted a comprehensive functional survey of the druggable mammalian epigenome, uncovering several elusive epigenetic drivers of treatment resistance in cutaneous squamous cell carcinoma (cSCC). This revealed *PAK1* as a previously unrecognized mediator of cSCC resistance in human and mouse SCC, offering new insights into a prognostic marker and therapeutic target of high clinical significance. Our findings establish RESTRICT-seq as a powerful tool for extending the applicability of pooled CRISPR screens to dynamic and previously intractable biological contexts.

## INTRODUCTION

CRISPR-Cas9 pooled screening has revolutionized functional genomics, uncovering many cancer cell vulnerabilities in nearly any cell type or context^1–6^. Over the past years, additional tools such as CRISPR interference (CRISPRi) and activation (CRISPRa) have expanded the precision of pooled screens^7,8^. However, despite their robustness, these systems lack DNA cleavage ability, making them insufficient for assessing absolute genetic necessity. Thus, knockout-based screens remain the gold standard for interrogating gene essentialities and dependencies at scale. In a typical CRISPR knockout screen, genetically perturbed cells compete for representation under a defined selective pressure (e.g., drug treatment) to determine gene function in that tailored contexts^9^. This largely relies on constitutively active Cas9 systems, which are burdened by off-target effects^8,10–12^, unintended cellular stress^13–18^, reduced cell fitness^11,19–22^, and stochastic gene expression changes that compound over time^22,23^. Consequently, constitutive Cas9 induction can severely distort clonal trajectories – which is detrimental for studying drug synergy, as therapeutic pressures can magnify cellular vulnerabilities and gRNA read count inequality^10,24^. Notably, even non-cleaving Cas9 variants such as CRISPRi/a exhibit negative fitness effects on cells, underscoring the broad disruptive impacts of nuclear-bound Cas9 beyond double-strand breaks (DSBs)^19,25^. Therefore, dropout screen outputs can often reflect the clonal progeny of a biased subset of initially transduced cells^10,22^, a major source of false-positive findings that obscure true biological signal and undermine screen fidelity^26^.

Several approaches, including clonal tracking of perturbed cells^13,27^, inducible Cas9 systems^28–31^, and transient delivery of Cas9–sgRNA complexes^19,32–34^ have been developed to mitigate clonal drift in pooled screens. Yet, none allow precise, repeated control of Cas9 nuclease activity throughout a screening experiment. These limitations hinder the ability to synchronize editing exclusively with drug treatment windows to prevent the accumulation of off-target effects and artefactual stress that drive clonal distortions in pooled screens. A few prior reports have demonstrated that titrating Cas9 nuclear concentration and residence time reduces off-target activity in a dose– and time-dependent manner^31–33^, offering a promising strategy to mitigate the unintended effects caused by prolonged Cas9 exposure. We hypothesized that restricting Cas9 catalytic activity and nuclear residence to biologically relevant time windows – particularly those aligned with drug exposure – would reduce artefactual clonal drift and improve signal fidelity in proliferation-based CRISPR screens. To investigate this, we developed the Repetitive Time-Restricted Editing followed by Sequencing (RESTRICT-seq) assay, a next-generation CRISPR screening method that strictly synchronizes Cas9 catalytic activity with biologically relevant time points. In contrast to conventional screens, RESTRICT-seq relies on the allosterically regulated Cas9 (arCas9) system, whose nuclear localization and catalytic induction are precisely controlled by ligand-receptor activation^35^. This enables repeated, pulsed editing at precise time windows, preserving clonal hierarchies in perturbed cells and improving the fidelity of gRNA dropout measurements over time.

We further benchmarked RESTRICT-seq across numerous screening contests, including conventional wtCas9 constitutive systems and traditional inducible protocols with both epigenome-wide libraries and kinome-wide pooled screen dataset^36^. As a test case, we used AZD4547, a selective inhibitor of fibroblast growth factor receptors (FGFRs), associated with high rates of resistance acquisition and currently under clinical investigation for use in combination therapy across multiple solid tumors, including squamous cell carcinoma^37,38^. Applying RESTRICT-seq to a skin cancer model of anti-FGFR therapy resistance^39^, we uncovered several novel epigenetic regulators critical for early-stage cancer resistance, including *PAK1* – a gene uniquely identified by RESTRICT-seq but missed by conventional screen approaches. Our findings reveal that maximizing synchronization between editing timing and drug exposure offers a critical advance in mitigating the pervasive confounders that limit current pooled screening methods. This work expands the experimental design space for pooled genetic screening in contexts where dynamic or proliferation-sensitive processes are being investigated.

## RESULTS

### arCas9 is an efficient gene editing system in skin cells

A critical feature that sets the allosterically-regulated Cas9 (arCas9) system apart is the lack of nuclear localization signal (NLS) and lack of nuclear export signal (NES). Instead, the arCas9 system relies solely on 4-OHT-mediated allosteric shift for movement in and out of the nucleus^35^. To benchmark the efficiency of this function, we first examined the baseline intracellular distribution of arCas9 by immunofluorescence staining of Human Embryonic Kidney (293T) cells expressing the pBLO1811-arCas9-noNLS-mCherry vector (293T-mCherry) prior to 4-OHT exposure (**Supplemental Figure 1A**). This revealed that arCas9 was primarily localized in the cytoplasm. Yet, in a small number of cells (42.55 ± 17.0%), arCas9 expression overlapped with DAPI staining, indicating low-level passive nuclear diffusion of arCas9, even in the absence of 4-OHT (**Supplemental Figure 1B)**. Subsequent immunofluorescent analysis of the mCherry protein, encoded via a T2A sequence downstream of the arCas9 gene, revealed even stronger overlap with DAPI (**Supplemental Figure 1B; Supplemental Figure 1G)**. Given that fetal bovine serum (FBS) contains trace amounts of estrogen^40^ and that phenol-red dye can act as a weak estrogen-like molecule that interferes with the estrogen receptor^41,42^, we hypothesized that FBS in phenol-red containing media contributed to low-levels of unspecific arCas9 nuclear translocation. To investigate this, we grew cells in phenol red-free culture media containing activated charcoal-stripped FBS, which depleted all lipophilic steroid molecules including estrogen (**Supplemental Figure 1C)**. We observed that 293T-mCherry cells cultured in charcoal-stripped phenol red free (CSPF) media displayed generally lower passive arCas9 nuclear localization (29.96 ± 8.70%) (**Supplemental Figure 1D; Supplemental Figure 1G**). Importantly, adding 4-OHT to media resulted in a robust increase in arCas9 nuclear translocation (99.5 ± 0.7%), irrespective of the media formulation (**Supplemental Figure 1H**). Moreover, we found that DMEM supplemented with unstripped FBS did not increase background gene editing in cells under vehicle treatment (OFF) when compared to scramble control 4-OHT (ON) treatment (**Supplemental Figure 2H-I**), indicating that low-level passive nuclear translocation does not significantly affect arCas9’s editing precision. Nevertheless, given that nuclear Cas9 can negatively impact cell fitness even when no specific gene is targeted^4,5,10,20,23,25,43^ and CSPF media was well tolerated by cutaneous SCC (cSCC) cells (**Supplemental Figure 1I**), we therefore used CSPF media in all subsequent experiments to further minimize passive arCas9 nuclear entry.

Next, to validate the efficacy of arCas9 in mammalian skin cells, we assessed its editing efficiency in various cell lines using fluorescent reporters and flow cytometric analysis (**Supplemental Figure 2A-E**). We used 293T cells, transfected to stably express both EGFP and arCas9-mCherry, to test arCas9’s ability to perturb the EGFP transgene^44^. In this experiment, cells expressing EGFP-targeting guide RNAs (sgEGFP) but treated with the ethanol (EtOH) vehicle control showed no significant reduction in EGFP-positive cells (76.8 ± 10.6%) compared to 4-OHT treated cells expressing a scramble gRNA (sgSCR; 79.0 ± 3.0%) (**Supplemental Figure 2G-H**). However, upon 4-OHT treatment, cells expressing sgEGFP showed a 4.4-fold decrease in EGFP-positive cells (22.57 ± 3.0%) compared to EtOH-treated sgEGFP cells and 3.4-fold relative to 4-OHT-treated sgSCR controls (**Supplemental Figure 2G-H**). These results confirmed arCas9’s inducibility for catalytic activation and efficient gene perturbation (**Supplemental Figure 2F**).

To refine arCas9 for use in mammalian skin cells, we optimized its catalytic induction in cutaneous keratinocytes, which are particularly difficult to genetically manipulate^45,46^. For these experiments, the arCas9-mCherry plasmid was subcloned into a lentiviral vector and stably transduced into N/TERT cells to generate arCas9-mCherry-expressing keratinocyte cell lines. We next targeted exon 1 of the integrin alpha 6 receptor (ITGα6) gene, a transmembrane receptor robustly expressed in skin basal keratinocytes. We found that EtOH-treated cells expressing ITGα6-targeting gRNA (sgITGα6) showed no significant decrease in ITGα6-positive cells (48.9 ± 30.8%) compared to 4-OHT-treated sgSCR controls (62.24 ± 8.9%) (**Supplemental Figure 2I**). However, sgITGα6-expressing cells displayed a significant reduction of ITGα6 expression (7.8 ± 4.5%) after 72 hours of exposure to 300 nM 4-OHT, a 4.4-fold and 3.4-fold decrease relative to EtOH-treated sgITGα6 cells and 4-OHT-treated sgSCR controls, respectively (**Supplemental Figure 2I**). These experiments confirmed the efficiency of the arCas9 system in skin cells and demonstrated that treating arCas9-expressing skin cells with 300nM 4-OHT for 72h was sufficient for significant decrease in target protein expression.

### Developing arCas9^SPARK^ to calibrate arCas9’s allosteric response

To rigorously investigate the kinetics and inducible precision of arCas9 nuclear translocation, we developed an arCas9 biosensor variant capable of real-time intracellular protein tracking in live skin cells (**Supplemental Figure 3**; **Figure 1A-D**). However, the complexity of arCas9, already modified to include the human estrogen receptor 1 (ESR1) ligand-binding domain (LBD) and designed for ligand-dependent allosteric regulation, posed significant challenges for further domain modifications^35^. We therefore created an *in silico* pipeline for enabling strategic insertions of additional functional domains into arCas9. This pipeline segmented arCas9 into six modular regions, allowing for customizable domain insertions and substitutions (**Supplemental Figure 3A-D**).

**Figure 1.**
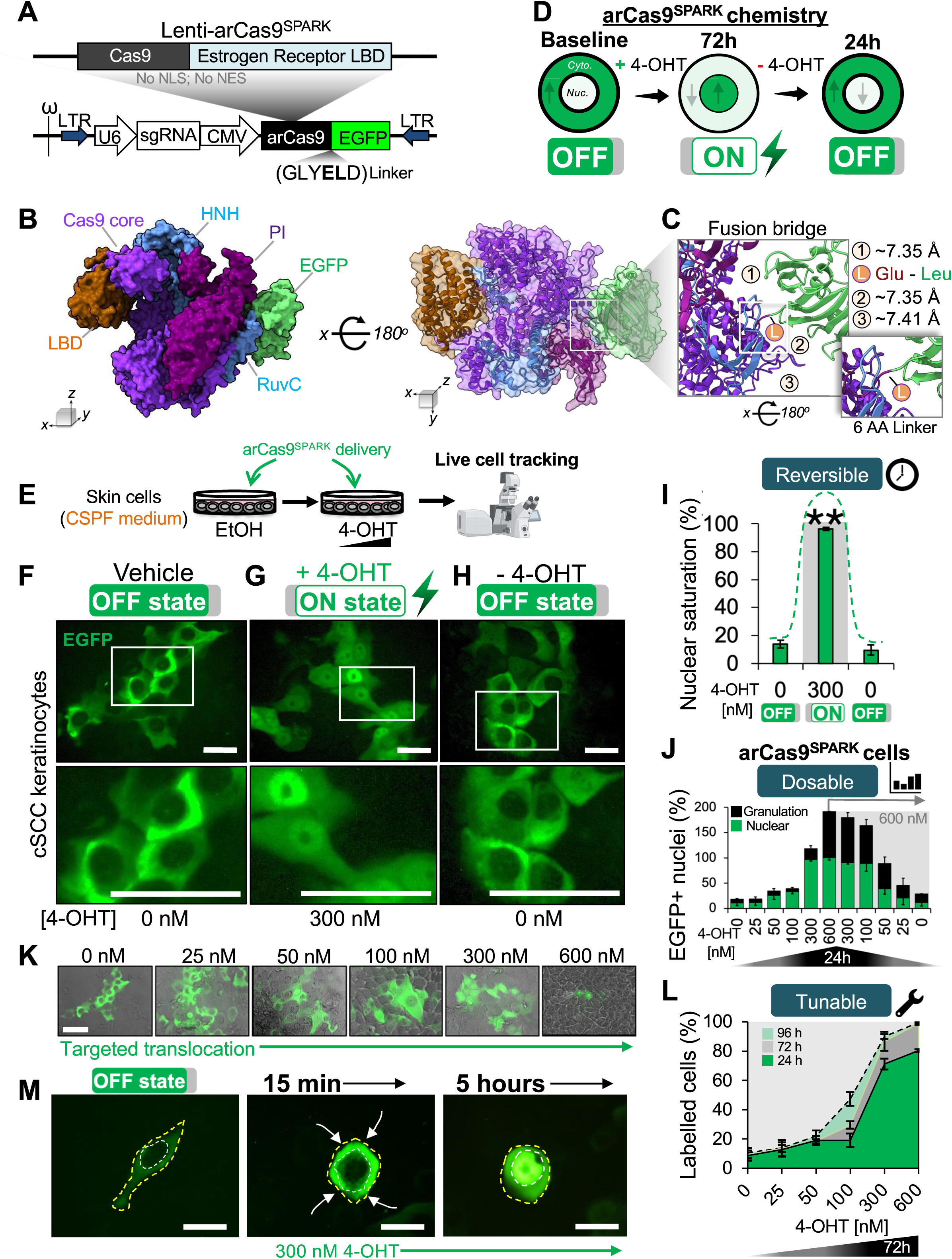
arCas9^SPARK^ is a ligand-responsive CRISPR biosensor tracking arCas9 translocation kinetics. (A) Structural map of the lenti-arCas9^SPARK^ plasmid. (B) 3D protein structure of arCas9^SPARK^, showing the spatial arrangement of its functional domains, as visualized in ChimeraX. (C) The arCas9^SPARK^ 6-amino acid (AA) linker (L) creates a fusion bridge between glutamic acid residue 1424 from the PAM interacting domain and leucine residue 1410 from the EGFP beta barrel, within the fusion protein. This configuration forms small cavities (1, 2, 3) which permit steric stability and flexibility, allowing for the dynamic conformational changes necessary for arCas9 function. (D) Theoretical overview of 4-OHT ligand-controlled arCas9-^SPARK^ nuclear oscillation during the ON (+4-OHT) and OFF (–4-OHT) activation states. Cytoplasm is the outer rim, while the nucleus is the inner rim. (E) Experimental outline of the arCas9-^SPARK^ nuclear oscillation (F) Representative images of live cell analysis showing arCas9-^SPARK^ subcellular localization in SCC-13 cells under vehicle treatment, (G) after 72h of 300 nM 4-OHT treatment for induced activation, (H) and 24h post-replacement of 4-OHT media with vehicle. The white box designates area zoomed in the bottom panel. *n* = 3 technical replicates from one representative cell line. Scale bar, 50μm. (I) Percentage of EGFP+ nuclei in arCas9-^SPARK^-expressing SCC-13 cells during the OFF state, after treatment with 300 nM 4-OHT, or after removal of 4-OHT. (J) Percentage of arCas9-^SPARK^-expressing cells with EGFP-saturated nuclei during treatment with various doses of 4-OHT for 24h (green bars). (Right) Percentage of EGFP-saturated nuclei 24h after 4-OHT removal in cells previously treated with 600 nM 4-OHT. Black bars depict the rate of cellular granulation. (K) Representative Phase contrast and EGFP overlaid images of arCas9-^SPARK^-expressing SCC-13 cells exposed to increasing doses of 4-OHT treatment. Scale bar, 50μm. (L) Percentage of arCas9-^SPARK^-expressing cells with nuclear EGFP expression after various 4-OHT treatments across 96h. (M) Representative images of intracellular EGFP localization tracking in arCas9-^SPARK^-expressing cells treated with 300 nM 4-OHT over short and medium time intervals (15 minutes to 5 hours). The yellow outline demarcates the cell membrane; the white outline depicts the nucleus. Arrows depict movement patterns of the EGFP signal towards the nucleus. Scale bar, 10μm. Bar graphs represent the mean ± SD of *n* = 3 technical replicates from one representative cell line in two independent experiments. Statistical significance was determined using a two-sided Student’s unpaired t-test (*: p < 0.05, **: p < 0.01, ***: p < 0.001, ****: p < 0.0001).

Using this strategy, we simulated various arCas9 construct designs to identify optimal sites for inserting an EGFP tag, which would facilitate tracking of arCas9’s intracellular movements (**Supplemental Figure 3B-C**). We found that vector position 3 (residue S241), containing the LBD, was particularly amenable to such insertions. We also considered insertions at the N-terminus (vector position 1; residue D-1) and C-terminus (vector position 7; residue V1629). However, the simulations revealed that placing EGFP near N-terminus residue S241 could interfere with ligand binding functions due to the LBD proximity (<20 Å) **(Supplemental Figure 3E-G)**. In contrast, inserting EGFP at the C-terminus maximized the distance from the LBD (>40 Å), resulting in fewer steric clashes (11.75 ± 3.2) **(Supplemental Figure 3G-J).** These findings suggested that the C-terminal region of arCas9 is more suitable for additional functional domain insertions, providing a clearer path for subsequent engineering efforts.

To verify the efficacy of our protein assembly pipeline, we inserted an EGFP protein at vector position 7 (**Supplemental Figure 3C-D, G**). This modification resulted in an enhanced variant engineered for real-time tracking and sensing of arCas9 protein allosteric response and kinetics (SPARK), offering capabilities that extend beyond those of a standard EGFP fusion protein (**Figure 1A-D**). The arCas9^SPARK^ design includes a short amino acid linker which generates three small external cavities (∼7.37 Å ± 0.03 Å), providing the necessary steric flexibility for the conformational shifts required during arCas9’s allosteric activation and deactivations (**Figure 1B-C**). These cavities ensure that the EGFP insertion does not hinder arCas9’s conformational changes, thus maintaining both allosteric functionality and the EGFP fluorescence. To evaluate this system in live cells, arCas9^SPARK^ was subcloned into a lentiviral vector (Lenti-arCas9^SPARK^) (**Figure 1A**), then used to study the dynamics of arCas9 nuclear translocation in live cSCC cells subject to varied 4-OHT exposures (**Figure 1E**). Initially, cells expressing arCas9^SPARK^ in its catalytically inactive OFF-state (EtOH only) showed EGFP fluorescence confined to the cytoplasm (**Figure 1F**), validating the design’s effectiveness in keeping arCas9^SPARK^ cytoplasmic when not activated. To assess whether EGFP fluorescence was preserved in the active ON-state, cells were treated with 300 nM 4-OHT for 72 hours. This treatment led to a pronounced shift of EGFP fluorescence into the nucleus (96.3% ± 1.24), with only faint signal remaining in the cytoplasm (**Figure 1G, I**), confirming that arCas9^SPARK^ retains both fluorescence and the ability to undergo the allosteric conformational change necessary for its nuclear translocation.

Given that the arCas9 system lacks a nuclear export signal (NES)^35^, we used arCas9^SPARK^ to assess whether arCas9 could exit the nucleus post-translocation. As 4-OHT induces nuclear import by binding to cytoplasmic ESR1-LBD, we anticipated that removing 4-OHT would revert this process, allowing arCas9 to return to the cytoplasm. To test this, 4-OHT was withdrawn from treated cells and replaced with ethanol (EtOH) vehicle-only media for 24 hours. This resulted in robust export of the EGFP signal from the nucleus (9.67% ± 3.4) back to the cytoplasm, validating the reversibility of arCas9 nuclear translocation upon ligand removal (**Figure 1H-I**). Importantly, when cells were subjected to a regimen of escalating followed by decreasing 4-OHT concentrations, arCas9^SPARK^ demonstrated that arCas9 retains the ability to undergo dose-dependent nuclear export, even after being previously exposed to concentrations as high as 600 nM (**Figure 1J**). Collectively, these experiments highlight arCas9^SPARK^ as a precise tool for real-time tracking of arCas9’s allosteric kinetics in live cells.

### Optimizing arCas9 inducibility with arCas9^SPARK^

To develop a robust methodology that ensures pooled screens maximize synchronization of editing exclusively with a desired therapeutic treatment being tested, we sought to further calibrate arCas9 inducibility. Leveraging the Lenti-arCas9^SPARK^ system, we explored a range of 4-OHT concentrations to find the balance between maximal nuclear import/export and minimal cellular stress (**Figure 1J**). We identified 300 nM as the optimal concentration that facilitated maximal arCas9 nuclear translocation while maintaining low levels of cellular granulation, indicative of reduced stress in skin keratinocyte^47^ (**Figure 1J-K**). In contrast, higher doses of 4-OHT such as 600 nM, increased translocation efficiency to 92.03 ± 4.41%, but at the cost of significant keratinocyte granulation and stress (**Figure 1J-K**). We next evaluated the stability of this reversible nuclear-cytoplasmic shuttling for applications in perturbation studies. Previous studies in skin cells expressing estrogen receptor-fused Cre recombinase (Cre-ER) have shown that low doses of tamoxifen, the prodrug of 4-OHT, can induce efficient gene deletion^48^. To evaluate whether arCas9^SPARK^ retains this sensitivity and temporal responsiveness, we treated cSCC cells with varying concentrations of 4-OHT over multiple time points. We observed a clear dose– and time-dependent increase in arCas9 nuclear localization, reaching near peak efficiency (71.13 ± 3.72%) as early as 24 hours post-treatment with ≥ 300 nM 4-OHT (**Figure 1L**).

A major challenge in long duration *in vitro* studies is the persistent and often hidden effects that the cell culture environment has on the experiment^49,50^, which is a source of artefactual noise in genetic perturbation studies^12,50–52^. Efforts to titrate Cas9 dosage have been shown to reduce such noise, as Cas9’s off-target profile is dose-dependent and the concentration of nuclear SpCas9 impacts its degree of spurious editing^23^. We thus reasoned that selective and immediate ON-OFF control of Cas9 catalytic activity could be crucial to avoid confounding effects from routine cell culture procedures and constitutive nuclear Cas9 activity. We next tested arCas9 responsiveness to brief 15–30-minute pulses of 4-OHT treatment in arCas9^SPARK^-expressing SCC cells. This revealed prompt nuclear translocation of arCas9 as early as 15 minutes (**Figure 1M**), validating arCas9’s suitability for applications involving short, repeated ON-OFF oscillations. Next, we examined arCas9 susceptibility to allosteric oversaturation following prolonged ligand exposure, which we suspected could lead to ligand dissociation from the LBD and spurious arCas9 relocation to the cytoplasm. Similar effects have been observed in CRISPRi-based systems, where dCas9 can gradually dissociate from its target loci over time, limiting its effectiveness in extended experiments^34,53,54^. Thus, we used arCas9^SPARK^ to establish dose-dependent parameters for long-term (≥ 14 days) arCas9 induction stability. Since arCas9^SPARK^ is cytoplasmic when unbound to the ligand, the cytoplasmic EGFP signal approximates the unbound receptor, while nuclear EGFP signal signifies the ligand-receptor complex (**Supplemental Figure 4A**). Together with the known ligand concentration and the duration of ligand exposure, these measurements enabled us to apply the law of mass action^55,56^. Using this approach, we found that extended 4-OHT treatment resulted in increased nuclear retention of arCas9, accompanied by a time-dependent decrease in its dissociation constant, with maximal nuclear occupancy achieved as early as 3 hours post-exposure (**Supplemental Figure 4 B, F**). These findings reveal the induction robustness of arCas9 for applications requiring sustained control over gene editing, an ideal feature for pooled screens. Conversely, the native arCas9 variant and the wild-type LentiCRISPRv2GFP system (wtCas9), where fluorophore expression is linked via a T2A sequence^57^, had signals distributed uniformly throughout the cell (**Supplemental Figure 4B-E, Supplemental Figure 1B**). Altogether, our findings confirmed the lack of allosteric oversaturation in the arCas9 system and emphasized the importance of CRISPR^SPARK^ engineering for precise tracking of arCas9’s nuclear-cytoplasmic dynamics.

Next, we aimed to investigate the perturbation efficiencies of arCas9 in long-term experiments. While arCas9^SPARK^ is effective as a biosensor, its large size (≥178 kDa), resulting from multiple functional domains, poses challenges for efficient delivery into hard-to-transduce skin keratinocytes. Furthermore, the EGFP fusion near its PAM-interacting domain could interfere with DNA recognition (**Figure 1B-C**). Therefore, although arCas9^SPARK^ is well-suited for studying precise protein translocation dynamics, we opted for the native arCas9 variant when conducting functional gene knockout experiments. To directly correlate arCas9^SPARK^ translocation with editing efficiency, we leveraged the dual role of 4-OHT, which both induces arCas9 nuclear translocation and activates its DNA cleavage function. Thus, nuclear localization of arCas9 serves as a direct indicator of its capacity for genome editing and can be used as a proxy for editing efficiency, consistent with first-order reaction kinetics. To test this, we established a baseline gene-editing efficiency for the native arCas9 variant in the OFF state, compared to the ON state 72 hours after exposure to 300 nM 4-OHT. Editing efficiency was assessed under ON or OFF conditions across multiple cell lines (**Supplemental Figure 2H-I, Supplemental Figure 5L-M**). Next, we measured nuclear translocation dynamics using arCas9^SPARK^ at matching time intervals (**Supplemental Figure 4B**). These measurements revealed a striking positive correlation (*r* >0.9) between nuclear translocation rates and editing efficiency, confirming that nuclear translocation reliably reflects arCas9 editing activity. To predict arCas9 editing outcomes based solely on nuclear translocation dynamics observed with arCas9^SPARK^, we developed a linear regression algorithm named arCas9 Modeling for Activity Prediction (arMAP) (**Supplemental Figure 4G**). Using arMAP, we found that optimal gene perturbation could be achieved as early as 3 hours after ligand exposure in arCas9-expressing cells (**Supplemental Figure 4I**). In contrast, NLS-containing wtCas9 showed consistently high nuclear localization and editing activity even before 4-OHT exposure (0h) (**Supplemental Figure 4C, E, J; Supplemental Figure 7O**), indicating that conventional NLS-tagged wtCas9 rapidly accumulates in the nucleus and exhibits greater background editing activity even without a gene-targeting gRNA (**Supplemental Figure 4J**). Together, these findings demonstrate that arCas9 enables tunable and sustainable long-term nuclear localization and editing activity in mammalian cells offering a robust platform for temporal precision in prolonged gene-editing experiments. We thus sought to harness arCas9 for temporally controlled pooled screening.

### Establishing high-risk cSCC mouse models amenable to arCas9 pooled screens

We next applied arCas9 pooled CRISPR screening to uncover epigenetic mechanisms driving resistance to FGFR inhibition in SCC. For this, we employed three distinct autochthonous mouse models of high-risk cSCC (HR-cSCC) by expressing driver mutations in specific skin epithelial compartments: DMBA-treated *K14-CreER, Rosa26-LSL-P53^fl/fl^; K14-CreER, Notch1^L/+^ Pik3ca^E545K/+^ p53^fl/+^;* and *K19-CreER*, *Pik3ca^E545K/+^ p53^R172H/+^ Notch1^fl/fl^*(**Supplemental Figure 5A**, **Supplemental Figure 6A**). Considering that metastatic cSCC predominantly affects elderly and immunocompromised individuals^58–60^, we modeled aggressive tumor progression by serially passaging autochthonous tumors in both aged immunocompetent C57BL/6 mice and young NOD scid gamma (NSG) mice, lacking a functional adaptive immune system (**Supplemental Figure 5B, Supplemental Figure 6A**). This approach resulted in highly tumorigenic autochthonous cSCC tumors that developed metastatic lung nodules, a pathological phenotype that replicated upon subsequent orthotopic tumor passages in both NSG and C57BL/6J mice (**Supplemental Figure 5E-H**). To confirm that the metastatic lung nodules were derived from the primary cSCC tumor, we performed immunohistochemistry analysis of basal epithelial marker keratin 14 (KRT14) and the basal stem cell marker P63, preferentially expressed in the skin. Both orthotopic tumors and lung nodules showed similarly high expression levels of KRT14 and P63, confirming epidermal defined origin (**Supplemental Figure 5E**). Interestingly, we observed two distinct growth patterns in the autochthonous tumors: Tumor type I appeared as depressed, poorly demarcated lesions with a central area of ulceration or indentation. The surrounding skin showed slight thickening or erythema, suggesting infiltrative growth into deeper tissues without significant outward protrusion. In contrast, tumor type II presented as raised, well-demarcated, keratotic mass protruding outward from the skin, with intact surrounding skin and no visible ulceration or signs of inflammation (**Supplemental Figure 5C, Supplemental Figure 6B-E**), reminiscent of human HR-cSCC lesions^61^. To validate that the tumors retained their aggressive characteristics throughout *in vivo* passaging, we analyzed them with H&E staining after each passage, looking for signs of malignancy such as hyperplasia and keratin pearls (**Supplemental 5D, Supplemental Figure 6H-K**). Our analysis revealed that tumor cells from DMBA-treated *K14-CreER Rosa26-LSL-P53^fl/fl^* mice had significantly higher tumorigenicity and metastatic potential (**Supplemental Figure 5H, Supplemental Figure 6F-G**). However, only tumors from *K14-CreER Notch1^L/+^ Pik3ca^E545K/+^ p53^fl/+^* mice maintained tumorigenic and metastatic potential in both NSG and C57BL/6J mice (**Supplemental Figure 5H, Supplemental Figure 6F-G**).

We next established stable keratinocyte cell lines from each of the three tumor types after two *in vivo* passages (iP2) (**Supplemental Figure 5B, Supplemental Figure 6I-K**). The MR041718 cell line, derived from *K14-CreER Notch1^L/+^ Pik3ca^E545K/+^ p53^fl/+^* mice tumors and the 392-1 cell line previously derived from DMBA-treated *K14-CreER Rosa26-LSL-P53^fl/fl^* tumors^39^ demonstrated continuous proliferation beyond 30 population doublings without signs of senescence or growth arrest. These cell lines also maintained stable viability in suspension growth assays and showed resistance to differentiation in the keratinocyte differentiation media (**Supplemental Figure 7B-I**). Importantly, while both cell lines retained tumorigenic potential (**Supplemental Figure 6G-J, Supplemental Figure 7J**), the 392-1 line exhibited significantly higher proliferation and migration rates in culture (**Supplemental Figure 6I**, **Supplemental Figure 7C-E**). Surprisingly, the 0088/0089 cell line, derived from *K19-CreER*, *Pik3ca^E545K/+^ p53^R172H/+^ Notch1^fl;/fl^* tumors, demonstrated the least proliferative potential in culture and was unable to form orthotopic tumors in mice (**Supplemental Figure 7A-D**). To verify that these cell lines retained their keratinocyte identity in culture, we co-stained them with antibodies against KRT19 and KRT14 after at least three passages in keratinocyte serum-free medium (**Supplemental Figure 5G**). All lines expressed high levels of KRT14 and maintained an epithelial-like cobblestone morphology (**Supplemental Figure 6I-L**). Notably, only the 392-1 cells co-expressed elevated levels of both KRT19 and KRT14, while 0088/0089 cells lacked KRT19 expression (**Supplemental Figure 6L**), suggesting the latter are epithelial but may have lost expression of this marker in culture. Next, we generated cell lines expressing either Lenti-arCas9 or Lenti-wtCas9 in both MR041718 and 392-1 cells (**Supplemental Figure 5J; Supplemental Figure 7K-M**) and validated their editing capabilities using the EGFP reporter assay^44^ (**Supplemental Figure 2B, Supplemental Figure 5J-M; Supplemental Figure 7N-O**). We observed that gRNA targeting EGFP led to a robust reduction in EGFP expression for both wtCas9 and arCas9 ON cells after 72 hours, relative to the scramble gRNA control (**Supplemental Figure 5L-M; Supplemental Figure 7O**). This validated that both arCas9 and wtCas9 systems could effectively perturb genes in mouse skin tumor cells. Notably, we observed generally higher editing efficiency with wtCas9 compared to arCas9. However, significant EGFP repression was equally detected in sgRNA targeting the scramble control, indicating greater off-target gene perturbation by wtCas9 (**Supplemental Figure 5L-M**).

### Conducting RESTRICT-seq pooled screening with translocation-optimized arCas9

Given the robust proliferation and migration rates of 392-1 cells in culture (**Supplemental Figure 5I; Supplemental Figure 7B-I**), we employed them for subsequent pooled CRISPR screening experiments with an epigenome-wide pooled CRISPR library (**Figure 2A**; **Supplemental Table 1**). To uncover transient epigenetic drivers of HR-cSCC resistance to anti-FGFR therapy, we leveraged arCas9 to confine editing to defined temporal windows that would maximize signal over experimental noise. In this approach, arCas9 remains in the deactivated OFF state during the initial pooled library transduction and puromycin selection phases but is activated to the ON state by 4-OHT exposure only during treatment with the anti-FGFR inhibitor AZD4547^62^. However, unlike standard inducible screening protocols, 4-OHT is removed at every passaging step, deactivating cleavage activity during passive cell culture procedures irrelevant to the biological question of the screen. This strategy strictly synchronizes gene editing with the therapeutic pressure, confining arCas9’s induction to critical time windows that maximize drug-gene interactions (DGIs) and dependencies (**Figure 2B; Supplemental Figure 8A**). We termed this method Repetitive Time-Restricted Editing followed by Sequencing (RESTRICT-seq) and hypothesized that it would significantly improve the accuracy and reliability of epigenome-wide pooled screen under AZD4547 treatment.

**Figure 2.**
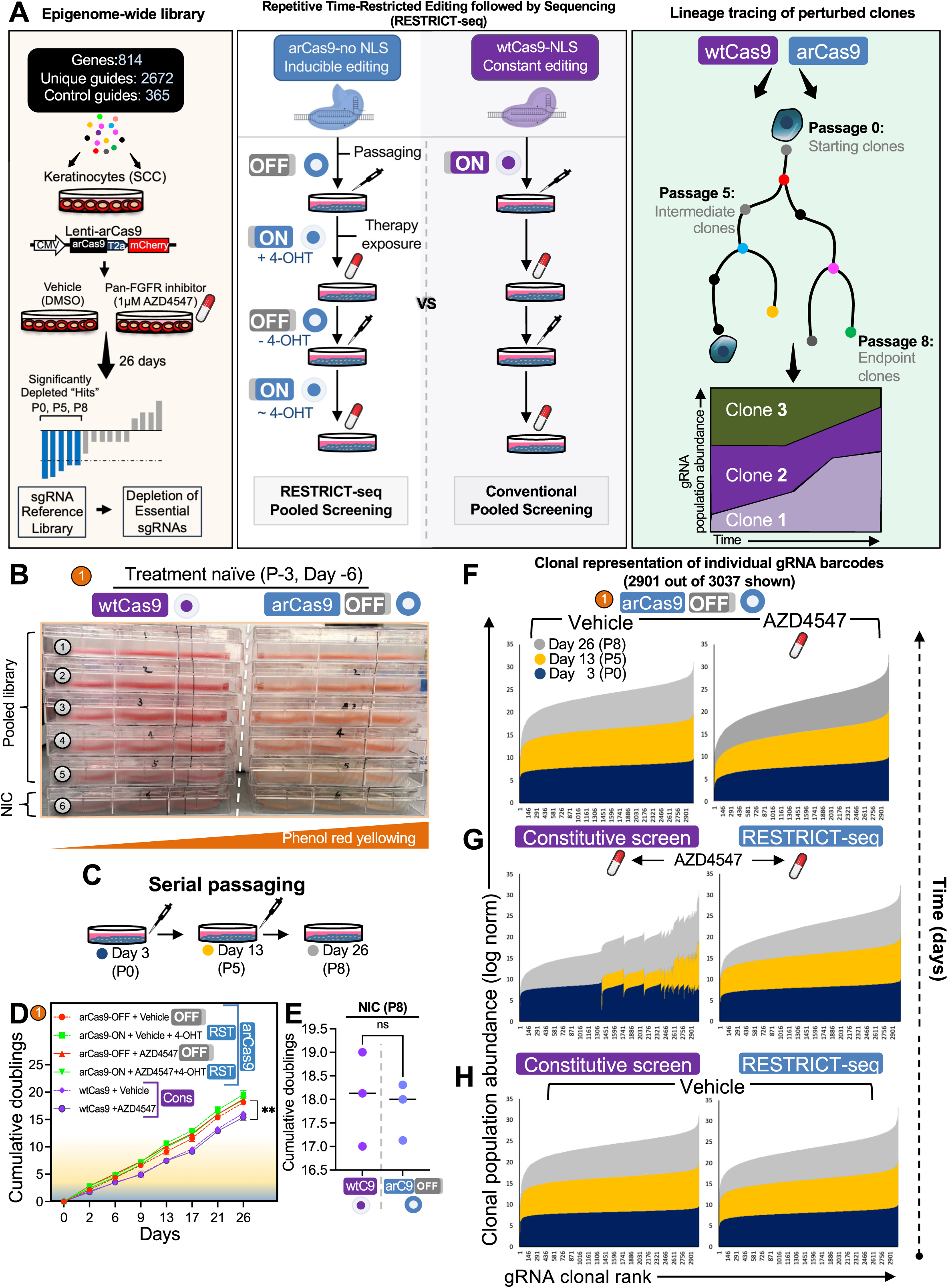
RESTRICT-seq maintains enhanced clonal fidelity in pooled screens. (A) Experimental outline of the RESTRICT-seq epigenome-wide pooled screen targeting all known epigenetic regulators expressed in mammalian skin. Cells are transduced with the pooled sgRNA library, and Cas9 activity is controlled intermittently, whereas it remains constitutively active in conventional screens. Population-level clonal tracing hierarchies are then reconstructed from perturbed cells to capture clonal evolution under screening conditions. (B) Assessment of cellular metabolic activity via phenol-red media yellowing in AZD4547-treatment naïve wtCas9 or arCas9-OFF 392-1 cells, either infected with the epigenome-wide library or serving as a no infection control (NIC) for 48 hours. *n* =30 technical replicates for each cell line. (C) Illustration of the serial passaging strategy used to monitor population doublings accumulated by perturbed cells over the course of the screen. (D) Cumulative population doublings of 392-1 cells under all pooled screening conditions over the course of 26 days post drug challenge. (E) Cumulative population doublings of non-infected control (NIC) 392-1 cells after CRISPR-OFF or conventional screens at P8. (F) Stacked bar graphs representing average clonal abundance of individual gRNA barcodes recovered from catalytically-inactive arCas9 screening (Deac, arCas9-OFF) in AZD4547-treated cells (right) and vehicle-treated cells (left) across three passage points over 26 days post-selection. *n* =3 technical replicates. (G) Average clonal abundance of individual gRNA barcodes recovered from RESTRICT-seq and constitutive wtCas9 screening in AZD4547-treated cells (top) and (H) treatment-naïve cells across three passage points over 26 days post drug challenge. *n* = 3 technical replicates from each representative cell line.

To benchmark RESTRICT-seq’s reliability, we conducted several additional CRISPR pooled screens in parallel (500x coverage): A) a standard inducible screen, where arCas9 was kept catalytically inactive during the spinfection and puromycin selection steps, but was constitutively activated after the AZD4547 treatment step. This is analogous to standard Cas9 inducible systems like *Cre/ER floxed* and doxycycline-inducible Cas9^63^. B) a catalytically-deactivated arCas9 screen in which cells were never exposed to 4-OHT throughout the duration of the screen. This is analogous to dead Cas9 systems^64^, which helped us determine leakiness in arCas9 screens. C) a constitutively active wild-type Cas9 (wtCas9) screen in which Cas9 remained catalytically active along the duration of the screen, including the spinfection steps (**Supplemental Figure 8A**). Next, we constructed a targeted epigenome-wide library comprising 2672 gRNAs, targeting all known epigenetic modifiers and associated regulators expressed in mouse skin. This library also included 197 essential genes to benchmark editing efficiency. Furthermore, to evaluate unspecific background activity, we incorporated an additional 365 non-targeting control (NTC) gRNAs, targeting intergenic regions or no gene at all (no target in the genome), as well as gRNAs targeting non-coding pseudogenes as negative controls. In total, the library targeted 814 genes, using 3637 individual gRNAs, with 4 separate gRNAs per gene for comprehensive coverage (**Figure 2A; Supplemental Table 1**). The library was subsequently transduced into arCas9-mCherry-expressing 392-1 cells and corresponding wtCas9-EGFP-expressing lines at a low multiplicity of infection (MOI) to ensure one gRNA per cell. Before puromycin growth selection (Day –6, passage –3), we assessed cell fitness and observed that library-transduced Lenti-arCas9 OFF cells showed normal metabolic activity, as evidenced by the acidification-induced yellowing of phenol-red media^65^ (**Figure 2D**). In contrast, library-transduced Lenti-wtCas9 cells exhibited reduced phenol-red yellowing, indicating editing-induced metabolic disruptions even before the start of the screen (**Figure 2B**). To evaluate how constitutive Cas9 induction affects screen performance, we quantified cumulative population doublings across multiple screening conditions (**Figure 2C**). Screens using sgRNA libraries in constitutively active wtCas9-expressing cells exhibited significantly lower cumulative population doublings compared to all other conditions, regardless of AZD4547 treatment exposure (**Figure 2D**). In contrast, no difference in population doublings was observed between NIC groups in wtCas9 and arCas9 cohorts (**Figure 2E**). These results align with previous findings that persistent DNA double-strand breaks (DSBs), which are continuously induced by wtCas9^11,14,66^, impair cell proliferative capacity by triggering cell cycle arrest and slowing division, resulting in a proliferative lag^5,67^ – a detrimental confounder in proliferation-based CRISPR screens. Altogether, these findings also corroborate ours (**Supplemental Figure 5L-M**) as well as prior reports of non-specific cleavage by wtCas9^52,67–71^. They highlight the critical importance of deactivating Cas9’s catalytic activity during non-essential phases of pooled screening to minimize unintended DNA cleavage, promote more synchronized cell cycle progression, and improve the overall fidelity of pooled CRISPR screens.

### RESTRICT-seq mitigates clonal population drift in pooled CRISPR screens

Our epigenome-wide pooled CRISPR screen spanned eight passages over 26 days, enabling temporal tracking of sgRNA abundance across long temporal scales. Each sgRNA vector delivered both a targeted genetic perturbation and a heritable barcode that could be used for tracing progeny^10^, allowing us to investigate cell clonal dynamics by quantifying the distribution of recovered gRNA counts in wtCas9– and arCas9-expressing cells (**Figure 2C**). As changes in gRNA representation reflect changes in population dynamics, they can provide a quantitative snapshot of the general distribution and relative abundance of clonal populations throughout the screen^72^. Previous studies have demonstrated that a major limiting factor in detecting gRNA-driven selection in wtCas9 is the high variance in gRNA counts between selection events^10^. To evaluate how different screening conditions affected these patterns, we measured cumulative clonal representation using the area under the abundance curve (AUC), which captures overall gRNA retention and dropout dynamics. We then quantified clonal drift as the standard deviation of gRNA abundance, representing the degree of dispersion and stochastic clonal expansion over time (**Supplemental Figure 8E-F**). In total, we monitored the abundance of 2,901 unique gRNAs, which enabled population-level reconstruction of clonal trajectories and revealed temporal clonal expansion or contraction patterns throughout the screen. Strikingly, RESTRICT-seq maintained a gradual and regular contraction of clonal abundance, with most clones retaining consistent representation and traceability from P0 to P8 (AUC = 0.90 ± 0.09) (**Figure 2G**; **Supplemental Figure 8E-F**). In contrast, the constitutively active wtCas9 screen exhibited abrupt dropout in cell clone representations, resulting in highly unstable and unpredictable clonal abundance pattens over time (AUC = 0.73 ± 0.38) (**Figure 2G; Supplemental Figure 8E-F**). Together, these results reveal that RESTRICT-seq better mitigates clonal population drift – even under therapeutic selection pressure – compared to constitutive Cas9 systems.

Next, to isolate the effects of AZD4547 from Cas9 endonuclease activity, we analyzed clonal abundance patterns in vehicle-treated cells. We found that clonal representation dynamics in vehicle-treated (DMSO) wtCas9 and RESTRICT-seq conditions were stable and comparable (**Figure 2H**; **Supplemental Figure 8E-F**), while AZD4547 treatment led to a larger reduction in clonal abundance across both editing systems (**Figure 2H-G**; **Supplemental Figure 8E-F**). To decouple the contribution of Cas9 endonuclease activity on clonal outcomes, we analyzed clonal distributions using the arCas9-OFF system. We found that AZD4547-treated arCas9-OFF screens exhibited negligible increase in clonal drift compared to their vehicle-treated counterparts (**Figure 2F**; **Supplemental Figure 8E-F**), reinforcing that Cas9 catalytic induction – rather than therapeutic pressure alone – primarily drives much of the observed artefactual drift in CRISPR screens. Strikingly, clonal abundance patterns of AZD4547-treated RESTRICT-seq screens (AUC = 0.90 ± 0.09) closely matched those of AZD4547-treated arCas9-OFF screens (AUC = 0.89 ± 0.03), underscoring that repeated catalytic deactivation under RESTRICT-seq is sufficient to mitigate undesired artefactual clonal drift that penalize synergistic CRISPR screens (**Figure 2F-G**; **Supplemental Figure 8E-F**). Given RESTRICT-seq’s superior ability to preserve clonal structures during prolonged drug exposure (>20 days), we sought to assess which confounding disruptions are best mitigated over the course of pooled CRISPR screens. We found that the RESTRICT-seq protocol captured 3-fold fewer disruptions, a 66% reduction in the total number of events included in constitutive wtCas9 screens (**Supplemental Figure 8B-D**). Notably, all disruptions captured by RESTRICT-seq were associated with biologically relevant selective pressures, such as cell refeeding with AZD4547 and clonal expansion under AZD4547 challenge (**Supplemental Figure 8A-D**). Altogether, this supports the conclusion that AZD4547 alone disrupts clonal composition independent of Cas9-mediated cleavage, but the induction of catalytically active Cas9 greatly exacerbates this effect. Our findings also revealed that precise control of Cas9 catalytic activity minimizes exposure to undesired clonal disruptions caused by routine cell culture steps, drastically reducing artefactual noise.

### RESTRICT-seq epigenome-wide screening identifies drug-specific vulnerabilities missed by conventional screens

Epigenetic alterations in keratinocytes are known to prime skin for neoplastic transformation and facilitate their progression from early-stage dysplasia to advanced cSCC^73–77^. To investigate whether similar epigenetic mechanisms contribute to resistance against anti-FGFR therapy, we confined RESTRICT-seq to defined AZD4547 treatment windows (**Figure 2A**) and compared gene dropouts relative to vehicle-treated controls (EtOH + DMSO) at three distinct time points across the 26-day screen: 1. Early (P0, Day 3), modeling pre-existing or rapidly acquired resistance. 2. Mid (P5, Day 13), reflecting emerging resistance. 3. Late (P8, Day 26) capturing stable resistance phenotypes^78^ (**Supplemental Figure 8A**). Next, to benchmark pooled editing efficiency, we compared RESTRICT-seq with conventional wtCas9 systems based on their capacity to deplete DepMap-defined common essential genes. Both systems yielded a similar total number of depleted genes irrespective of significance thresholds (160 for wtCas9 vs. 149 for RESTRICT-seq; 92.3% overlap, (**Supplemental Fig. 10C–D**). However, wtCas9 screens produced more unique common essential gene dropouts (56 vs. 39 unique to RESTRICT-seq; **Supplemental Fig. 10E**) but fewer depletions outside the common essential set (**Supplemental Fig. 9D, F**). Pathway-level analyses further revealed that wtCas9 screening identified a greater number of KEGG-enriched pathways prior to treatment but produced significantly fewer drug-dependent pathways than RESTRICT-seq (**Supplemental Fig. 10A–B**). Critically, RESTRICT-seq uncovered several novel drug-dependent genes, including *KAT2B* and the p21-activated kinase 1 (*PAK1*); annotated as *PKN1* in the Broad Institute CRISPR library (**Figure 3A; Supplemental Figure 9A-C**). Notably, PAK1 was not detected as significant in AZD4547-treated wtCas9 screens or across all arCas9-OFF screens (**Supplemental Figure 9D-F; Supplemental Figure 10F-G**), confirming its selective dependency on arCas9 activation exclusively during AZD4547 drug exposure. Conversely, KAT2B was depleted in both arCas9-ON and OFF conditions after drug exposure – suggesting a false-positive dropout event unrelated to actual editing activity (**Figure 3A**; **Supplemental Figure 10G).** This type of misclassification was undetectable using conventional wtCas9 screening, since an intrinsic OFF state could not be achieved within that platform. Instead, a parallel negative control using an entirely separate system would have been required, substantially increasing confounding variables to the experiment. Indeed, wtCas9 screens primarily identified common essential genes such as *TADA2A* and *IKZF1* as top dropouts (**Figure 3A**; **Supplemental Figure 9D, F**), reflecting limited specificity toward AZD4547 treatment and a higher prevalence of non-selective gene depletions. Remarkably, only arCas9 enabled clear disentanglement of non-specific or artifactual dropout events from genuine on-target effects within a single, unified pooled screening study. This highlights a tradeoff between breadth of editing activity and pathway-level specificity when using wtCas9 versus RESTRICT-seq. These results suggest that temporal gating in RESTRICT-seq preserves the ability to efficiently target genes in pooled screens but also yields a more refined dropout profile by minimizing contributions from non-specific or off-pathway hits. Altogether, these findings indicate that while wtCas9 better captures broad genetic essentialities, temporally gated editing with RESTRICT-seq enhances both sensitivity and specificity for drug-specific, context-dependent essentialities that are often obscured by constitutive editing and cumulative drift in conventional screens.

**Figure 3.**
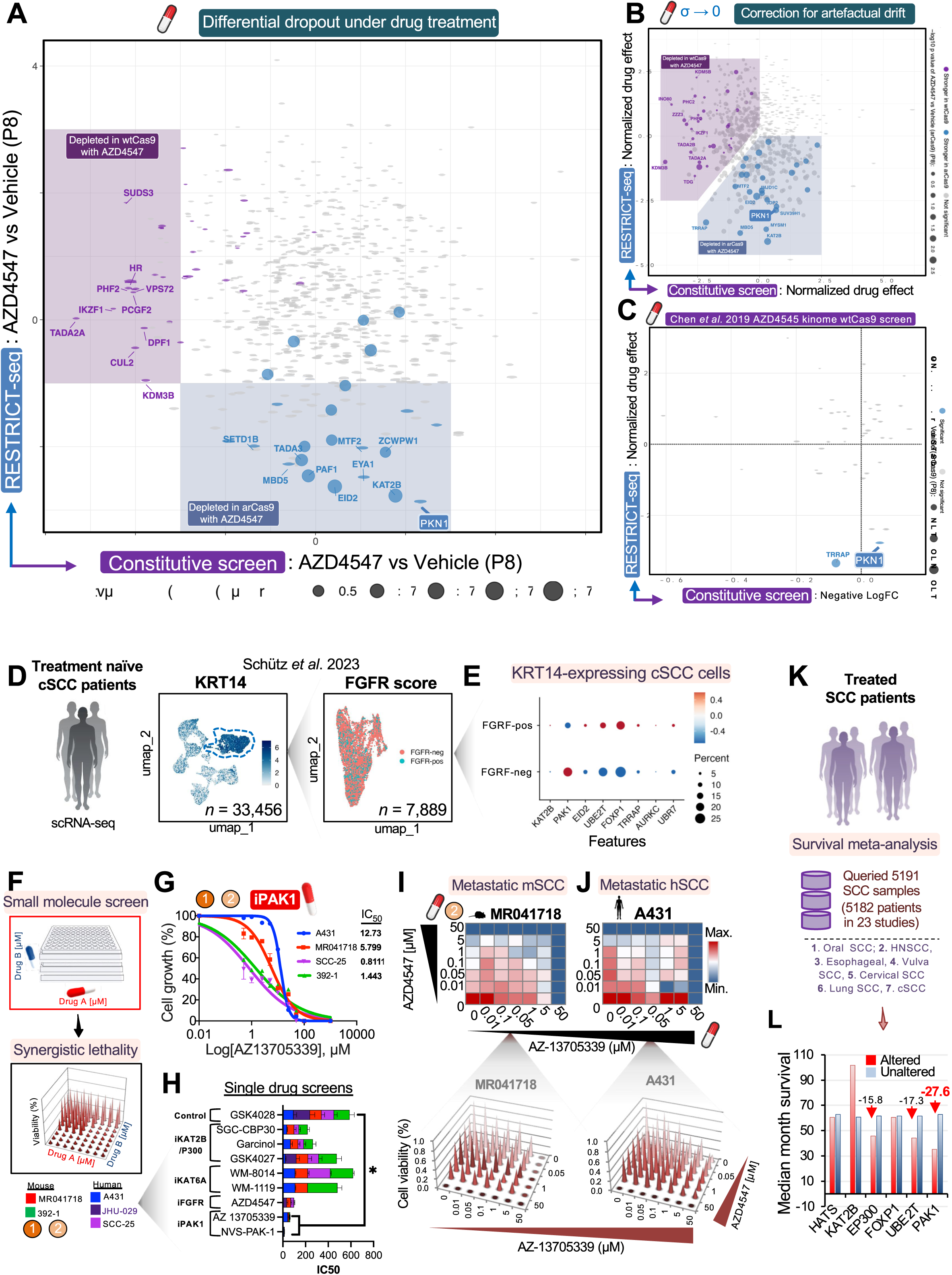
Multi-tiered normalization in RESTRICT-seq identifies PAK1 as an FGFR-specific SCC dependency. (A) Comparative Robust Rank Aggregation (RRA) analysis of depleted genes from AZD4547-treated RESTRICT-seq vs constitutive wtCas9 CRISPR screens at P8, relative to vehicle-treated controls. (B) RRA analysis depicting normalized drug effect in wtCas9 and arCas9-ON CRISPR screens. (C) Comparison of the normalized drug effect obtained from RESTRICT-seq with results from a previously published AZD4547-focused constitutive wtCas9 kinome-wide screen. Colored points indicate significant hits. (D) Retrospective single-cell RNA sequencing (scRNA-seq) analysis of FGFR expression levels in KRT14-expressing cells from treatment-naïve cSCC tumors. *n* = 3 biological replicates. (E) Expression levels of top significantly depleted target genes from RESTRICT-seq AZD4547 screen in FGFR-positive and FGFR-negative K14^+^ cells from treatment-naïve cSCC tumors. (F) Schematic overview of the small molecule inhibitor screen analysis of synergistic lethality. (G) 10-point dose-response inhibition screen and IC50 determinations for the PAK1 inhibitor AZ13705339 in human SCC cell lines, as well as 392-1 and MR041718 tumorigenic mouse cSCC cell lines. *n* = 3 technical replicates. (H) Average IC50 values (µM) for selective inhibitors targeting the top significantly depleted genes identified by RESTRICT-seq (i.e., KAT2B/EP300, KAT6A, PAK1), in mouse (MR041718, 392-1) and human SCC lines (A431, JHU-029, SCC-25). Assay included GSK4028 as enantiomeric negative control and AZD4547 as positive control. *n* = 3 technical replicates from three separate experiments. Statistical significance was determined using a one-way ANOVA followed by Dunnett’s test for multiple comparisons (*: p < 0.05, **: p < 0.01, ***: p < 0.001, ****: p < 0.0001). (I) Synergistic lethality matrix showing cooperative effects of AZD4547 with AZ13705339 in metastatic tumor-derived mouse (MR041718) and (J) human (A431) cSCC cells. (K) Schematic overview of the SCC human survival meta-analysis encompassing retrospective sequencing data from over 5,000 SCC tumor patients. (L) Overall median months survival of SCC patients with wild-type (unaltered) or altered alleles encoding top significantly depleted target genes identified by RESTRICT-seq. Red arrows depict relative difference in overall survival months among patients with altered alleles.

To further isolate gene dependencies exclusively synergistic with AZD4547 treatment, we designed a multi-tier normalization strategy to control for confounding factors arising from passaging artifacts and Cas9-mediated genotoxicities. We started by using differential dropout genes significantly depleted in AZD4547-treated arCas9-ON screens versus vehicle control at passage 8 (P8) (**Supplemental Figure 9A-C**). Next, to remove gene dropouts attributable to editing but independent of AZD4547 action, we subtracted gene hits differentially depleted in vehicle-treated arCas9-ON versus arCas9-OFF screens at P8 (**Supplemental Figure 9G**). Finally, to correct for editing-independent artefactual clonal drift arising from repeated passaging, we subtracted gene dropouts that emerged in vehicle-treated arCas9-OFF cells between baseline (P0) and P8 (**Supplemental Figure 10F**). Together, these stepwise normalization steps eliminated several major sources of confounding events common in synergistic CRISPR screen analyses and resulted in normalized drug effect that reliability reflected DGIs uniquely attributable to AZD4547 treatment (**Figure 3B**). For the constitutive wtCas9 system – in which Cas9 activity cannot be temporally deactivated – normalized drug effect was determined by comparing AZD4547-treated versus vehicle-treated (DMSO) samples at P8, followed by subtraction of gene depletions observed in vehicle-only conditions between P8 and P0 **(Figure 3B**). Importantly, *PAK1* was not detected as a significant dropout in any of the wtCas9 or arCas9-OFF screens (**Supplemental Figure 9F-G; Supplemental Figure 10F-G**) but was exclusively identified by RESTRICT-seq when both arCas9 editing and AZD4547-treatment were synchronized (**Figure 3A-B; Supplemental Figure 9G**). To determine whether other CRISPR studies detected *PAK1* as a FGFR dependency, we analyzed a previously published kinome-wide CRISPR dropout screen that also investigated AZD4547 treatment with the wtCas9 system^36^. While several essential gene dropouts overlapped between RESTRICT-seq and the kinome screen, *PAK1* was entirely undetected as a significant hit in the latter but remained exclusively identified by RESTRICT-seq (**Figure 3C**). This demonstrates RESTRICT-seq’s ability to disentangle drug-specific effects from background noise introduced by Cas9 activity and culture-induced stress, enabling selective detection of synergistic dependencies consistently missed by conventional CRISPR screening approaches.

### PAK1 inhibition abrogates SCC cell resistance to AZD4547 treatment

As a serine/threonine kinase, PAK1 regulates mitotic progression, and plays a pivotal role in diverse processes such as cell motility, invasion, and therapy resistance, through histone phosphorylation and modulation of tyrosine kinase receptor signaling, including EGFR^79–82^. Interestingly, while PAK1 dropout was identified as the most significantly depleted gene by P8, it was similarly detected at the earliest time point (P0, Day3) of the RESTRICT-seq AZD4547 screen, indicating a potential role in the cell’s predisposition to treatment resistance even before AZD4547 exposure (**Figure 3; Supplemental Figure 9A-C**). To investigate this, we analyzed single-cell RNA sequencing (scRNA-seq) data from treatment-naive HR-cSCC tumors^83^ and grouped K14^+^ cells based on cumulative levels of *FGFR1-4* expression (**Figure 3D**). Next, we examined expression patterns of top dropout genes uniquely identified by RESTRICT-seq, analyzing their levels at both early and late timepoints in FGFR-expressing tumor cells. Among these, *FOXP1* and *UBE2T* were most widely expressed in *FGFR*-positive tumor cells (> 10%), while the false-positive hit *KAT2B* was virtually undetectable in this cell group (< 1% of cells) (**Figure 3E**). Strikingly, *PAK1* was upregulated robustly (>20% of cells), and exclusively, in *FGFR*-negative tumor cells (**Figure 3E**). Together, this suggests that PAK1 functions as an early adaptive program abetting treatment-naïve SCC tumor cells for FGFR-independent survival.

To test this, we conducted a 10-point small molecule drug inhibition screen targeting PAK1 in five various human and mouse SCC cell lines. Remarkably, PAK1 inhibition consistently, and independently, reduced cell viability across both human and mouse SCC cell lines when compared to an enantiomeric control, a result validated using two structurally distinct PAK1 inhibitors (**Figure 3G; Supplemental Figure 11G**). Next, we performed combinatorial dose-response assays with the small molecule inhibitors to investigate the therapeutic synergy of co-targeting PAK1 and FGFR in metastatic HR-SCC cells. This revealed strong, dose-dependent synergy between PAK1 inhibition and AZD4547 treatment, resulting in a 20–40% reduction in cell viability relative to single-agent treatments alone – even at concentrations as low as 50 nM across human (A431, SCC-25) and mouse (392-1, MR041718) SCC cell lines (**Figure 3I-J, Supplemental Figure 12B**). These results demonstrate that co-inhibition of PAK1 and FGFR synergistically suppresses HR-SCC cell survival, supporting recent findings of PAK1’s involvement in therapeutic cross-resistance in prostate cancer^79^.

### PAK1 tumor amplification defines a prognostic axis of treatment response in human SCC

In addition to tyrosine kinase receptors and histones, PAK1 phosphorylates histone acetyl transferases (HATs), which form the PAK1/CREB phosphoacetylation axis critical for SCC cell survival^81,82,84,85^. Since HATs also emerged as top dropout genes identified by RESTRICT-seq under AZD4547 treatment (**Figure 3A-B**; **Supplemental Figure 9G**), we chose to equally interrogate whether targeting HATs, specifically KAT6A and CBP/P300, could likewise sensitize SCC cells to AZD4547 therapy. We found that garcinol – a dual inhibitor of KAT2B and CBP/P300^86^ – only modestly reduced IC_50_ values compared to enantiomer control (IC_50_ = 66.2 μM ± 11.9 vs. 117.6 ± 22.8 μM; p < 0.01; **Supplemental Figure 11C, H**). Moreover, selective inhibition of KAT2B alone using GSK-4027 did not meaningfully affect SCC cell viability (IC_50_ = 94.99 μM ± 48.3; **Figure 3H; Supplemental Figure 11A-B, H**), suggesting that garcinol’s modest effect against SCC cells was primarily attributable to CBP/P300 inhibition. To test this directly, we treated SCC cells with the CBP/P300-selective inhibitor SGC-CBP30, which significantly impaired SCC cells viability and yielded lower IC_50_ values (IC_50_ = 54.5 μM ± 19.2; p < 0.005) compared to enantiomeric control (**Figure 3H; Supplemental Figure 11D, H**). However, when combined with AZD4547, SGC-CBP30 produced largely additive rather than synergistic effects (**Supplemental Figure 12A, C-D**), indicating that while P300 inhibition alone is modestly cytotoxic to SCC cells, it does not strongly enhance the therapeutic efficacy of FGFR inhibition in this context. We next similarly evaluated whether KAT6A may serve as a synergistic partner with FGFR inhibition. Surprisingly, direct inhibition of KAT6A with either WM-1119 or WM-8014 alone did not significantly affect SCC cell survival (**Figure 3H**; **Supplemental Figure 11E-F**). However, when combined with AZD4547, WM-8014 caused marked cytotoxicity across SCC cell lines, with clear evidence of synergy in human A431 cells (**Supplemental Figure 12 E–F**). Together, these findings indicate that the efficacy of co-targeting HATs is more context-dependent, warranting further mechanistic studies to delineate their contribution to anti-FGFR therapy resistance.

Therefore, to evaluate the broader clinical relevance of our findings, we tested whether RESTRICT-seq–identified genes carried prognostic value in SCC. A meta-analysis of 5,182 patients spanning seven SCC subtypes across 23 treatment centers revealed that, despite chemotherapy, patients with *PAK1* amplifications had a 27.6-month lower median survival than those without alterations. Likewise, *EP300* amplifications were associated with reduced median survival (15.8 vs. 61.6 months; **Figure 3K–L**). These clinical trends are consistent with our drug inhibition results, which showed that selective P300 targeting modestly, but significantly, reduced IC_50_ values, though not as potent as PAK1 inhibition (**Supplemental Figure 11C-D, G**). Next, we analyzed clinical outcomes of patients harboring genomic alterations in a broader panel of HATs, including *EID2*, *KAT6A*, *KAT2B*, and *CREBBP*/*EP300*, as well as other top candidates identified by RESTRICT-seq, such as *FOXP1* and *UBE2T*. While *FOXP1* alterations did not have a considerable effect on patient survival outcome (∼ 1 month reduction), *UBE2T* alterations were associated with a striking 17.3-month reduction in survival (**Figure 3K**). In contrast, *KAT2B* alterations correlated with a 41-month increase in survival, corroborating results from our small molecule screen showing the minimal impact of KAT2B inhibition on SCC viability (**Figure 3H**; **Supplemental Figure 11B, H**). Interestingly, patients with at least one genomic alteration in the set of HAT genes only experienced a modest 9.9-month reduction in survival, reinforcing the notion that the therapeutic efficacy of many epigenetic targets is likely SCC subtype dependent. Altogether, our results establish a strong concordance between RESTRICT-seq identified vulnerabilities and patient outcomes, highlighting both the therapeutic potential and prognostic value of targets such as *PAK1* and *P300*.

## DISCUSSION

Collectively, our data demonstrate that conventional drug-synergistic pooled CRISPR screening approaches are constrained by substantial technical limitations; most notably, clonal distortions resulting from premature and cumulative, unrestricted Cas9-mediated editing. These issues are further exacerbated by routine cell culture manipulations, which induce cellular stress and compound variability in gRNA representation, ultimately obscuring true gene–phenotype relationships in dynamic biological contexts. Our study introduces a new, temporally gated screening strategy, RESTRICT-seq (Repetitive Time-Restricted Editing followed by Sequencing), designed to overcome these challenges. By precisely confining Cas9 catalytic activity to strict temporal windows through ligand-controlled allosterically regulated Cas9 (arCas9) activation, RESTRICT-seq enables controlled and cyclic induction of editing, which significantly mitigates undesired fitness penalties that accumulate over time in traditional screens.

This new CRISPR screening approach uncoupled non-specific artefactual dropout events from true on-target effects by strictly synchronizing gene editing with therapeutic pressure while deactivating Cas9 during passive culture steps. As a result, RESTRICT-seq yielded markedly improved consistency in gene dropout signatures and delivered more reliable gene hits that closely aligned with clinical patient outcomes. A pivotal discovery made with RESTRICT-seq was the identification of *PAK1* as a previously unrecognized mediator of cutaneous squamous cell carcinoma (cSCC) resistance to anti-FGFR therapy, presenting a prognostic marker and therapeutic target of high clinical significance. Notably, *PAK1* depletion emerged at the earliest treatment stage (P0) that validated using multiple small-molecule inhibitors across human and murine SCC models and showed strong dose-dependent synergy with AZD4547 compared to single-agent treatments alone. At a clinical level, meta-analysis of patient tumors showed that *PAK1* amplifications were associated with lower median patient survival (27.6-month reduction) than patients without amplifications. Strikingly, *PAK1* was consistently missed as a synergistic hit by conventional wtCas9 screens and arCas9-OFF controls.

Moreover, RESTRICT-seq offers several distinct advantages as compared to standard inducible systems, including compatibility with titratable repeated activation cycles and quick on/off switching. For example, while Tet systems regulate Cas9 abundance on timescales of days through *de novo* translation^87^, arCas9 nuclear translocation and inducibility occurs on the scale of minutes to hours without requiring new transcriptional cycles. This pulsing capability and rapid toggling make RESTRICT-seq ideal for multi-phase screens requiring interrogation of gene function during varied adaptive stages. Moreover, the inclusion of a catalytically deactivated arCas9-OFF screen served as a critical internal control, demonstrating that Cas9 catalytic induction, rather than therapeutic pressure alone, primarily drives much of the observed artefactual drift in pooled screens. Therefore, a sophisticated multi-tier normalization strategy was employed to control for confounding factors like passaging artifacts and Cas9-mediated genotoxicities, ensuring identified drug-gene interactions were uniquely attributable to AZD4547 treatment. In contrast, wtCas9 systems consistently exhibited lower cumulative population doublings and greater background activity, reflecting persistent DNA double-strand breaks and cleavage at undesired time windows. Because constitutive Cas9 induction can severely distort clonal trajectories and mask context-specific dependencies, its proper assessment is detrimental for studies investigating drug synergy. Perturbation screens should precisely control Cas9 activity to avoid confounding effects from routine cell culture procedures and constitutive nuclear Cas9 activity.

This study provides evidence that the confounders generated by routine cell culture handling and passive drug exposure during proliferation-based pooled screens can be mitigated by the RESTRICT-seq screening approach. Despite this significant advancement, RESTRICT-seq, like any tool, has constraints. For instance, a notable limitation of the arCas9 system for inducible pooled screens is its inherent complexity, particularly its 4-OHT ligand-dependent regulation and strict dependency on the optimal ligand dosing, which necessitated rigorous benchmarking. This need to characterize the system’s inducible activity was critical for skin cells and motivated the development of the arCas9^SPARK^ variant, an EGFP fusion biosensor designed for real-time intracellular allosteric tracking of arCas9. The arCas9^SPARK^ system was indispensable in elucidating arCas9’s nuclear kinetics, demonstrating reversible translocation, rapid on/off switching, and sustained dose-dependent induction stability. Such meticulous calibration requirements may pose challenges when applied to more sensitive primary cell types that may not tolerate prolonged calibration procedures or extended ligand exposures. Despite these considerations, RESTRICT-seq introduces a next-generation paradigm for pooled CRISPR screening by achieving higher signal-to-noise ratios than conventional tools and isolating clinically meaningful, drug-induced dependencies. Looking ahead, RESTRICT-seq’s compatibility extends beyond arCas9 and is amenable to other inducible platforms such as split-Cas9 systems^28,88,89^. This establishes a robust foundation for expanding pooled CRISPR screens to other dynamic biological processes and disease models where transient or proliferation-sensitive events are critical.

## MATERIALS AND METHODS

### Cell lines and culture conditions

Human embryonic kidney (HEK293T) and A431 cells were maintained in Dulbecco’s Modified Eagle Medium (DMEM, ThermoFischer Scientific) supplemented with 10% fetal bovine serum (FBS) or Bovine Calf Serum (BCS; ThermoFischer Scientific, SH3007203) and 1% Penicillin/Streptomycin (P/S; ThermoFischer Scientific, 15190122). Select experiments were performed in phenol red-free RPMI (ThermoFischer Scientific, 11835030) supplemented with 10% charcoal-stripped FBS (CSPF media) to eliminate steroidal ligands and minimize estrogenic interference. SCC-13, MR041718, 0088/0089 and N/TERT-1^90^ were cultivated in Keratinocyte Serum-Free Media (KSFM; ThermoFischer Scientific, 17005042) supplemented with 0.4M CaCl2, 0.25 ng/ml Epidermal Growth Factor (Life Technologies, 10450-013), 1/2 vial Bovine Pituitary Extract (Life Technologies, 13028014) and 1% P/S. JHU-029^91^ and 392-1^39^ cells were cultivated in RPMI 1640 media (ThermoFischer Scientific, 11875119) supplemented with 10% FBS or BCS and 1% P/S. SCC-25^92^ were cultured in DMEM/F-12 media (ThermoFisher Scientific, 11320082) supplemented with 10% FBS or BCS and 1% P/S. For differentiation-resistant culture conditions, KSFM was supplemented with 5% FBS and 1 mM CaCl₂ to induce spontaneous keratinocyte differentiation blockade in select assays as previously described^47^. All cell lines were cultured at 37°C in 5% CO₂ and passaged at ∼70% confluence. For clonal expansion and maintenance, antibiotic selection was performed with 1 µg/mL puromycin (Gibco) for stable cell line generation. Cell line identity was authenticated using STR profiling (ATCC, 135-XV). Expression of epithelial markers KRT14, KRT19, and p63 was confirmed by immunofluorescence and hyperplasia was confirmed by histological staining for keratin pearls.

### *In silico* structural modeling

Candidate designs were generated by assembling EGFP in various predefined plasmid insertion sites within the arCas9 coding sequence. Three-dimensional structural predictions were carried out by imputing desired sequences amino acid sequences using Phyre^2^ and AlphaFold3^93,94^. Each arCas9-EGFP fusion variant was assessed in ChimeraX v1.8 for predicted favorable spatial orientation relative to PAM binding and catalytic domains. For selected high-confidence designs, the corresponding nucleotide sequences were codon-optimized and subcloned into third-generation lentiviral backbone and tested into relevant skin SCC lines for live-cell imaging and functional validation of ligand-dependent nuclear translocation.

### Plasmid constructs and stable cell line generation

Stable fluorescent The Lenti-arCas9-mCherry plasmid was constructed by digesting the pLL3.7-EGFP vector^95^ with AgeI and EcoRI to create the insertion site, using the remaining plasmid backbone as the recipient vector. The pBLO1811_arCas9_noNLS-mCherry plasmid was equally digested with AgeI and EcoRI, and the arCas9 t2a mCherry fragment was subcloned into the AgeI/EcoRI-digested pLL3.7-EGFP backbone by sticky-end ligation. The optimal arCas9-EGFP fusion construct (arCas9-^SPARK^) was similarly generated by digesting pBLO1811_arCas9_noNLS-mCherry plasmid at Age1and BamH1 sites and subcloning the arCas9 fragment lacking mCherry directly upstream of the pLL3.7 vector EGFP sequence, creating a fusion protein. LentiGuide-puro vectors^96^ were clones using standard protocols^96^ EGFP targeting gRNA1: TCTCAGCACAGCATGGG, gRNA 2: CCTTGTCTCTAATAGAGG; and hITGA6 gRNA: GGAGCTCCGTATGATGACTT based on ChopChop^97^. Stable cell lines were established by transfecting HEK293T cells using pCMV-VSV-G and pCMV-dR8.2 dvpr packaging plasmids by the calcium-phosphate method (CalPhos Mammalian Transfection Kit, Takara, 631312) for virus production. Cells were fed with DMEM supplemented with 20% FBS and 1% P/S. Viral supernatants were collected 48– and 72-hours post-transfection, pooled and filtered through a 0.45 μm PVDF filter, then concentrated via ultracentrifugation. Target cells were spin-infected at 900 x g for 1 hour at 25°C in the presence of polybrene (0.8ug/mL Sigma), and the viral supernatant was then replaced with fresh growth media. Selection was started 72h later using puromycin (1 μg/ml) or through repeated rounds of flow cytometry sorting based on respective fluorescent transgene expression. The pBLO1811_arC9_noNLS_human was a gift from David Savage (Addgene plasmid # 74494). LentiCRISPRv2GFP was a gift from David Feldser^57^(Addgene plasmid # 82416). pLL3.7 was a gift from Luk Parijs (Addgene plasmid # 11795). LentiGuide-Puro was a gift from Feng Zhang (Addgene plasmid # 52963). PCMV-VSV-G was a gift from Bob Weinberg (Addgene plasmid # 8454). pCMV-dR8.2 dvpr was a gift from Bob Weinberg (Addgene plasmid # 8455).

### Translocation assays and arCas9-^SPARK^ biosensor modeling

Cells were treated with 4-hydroxytamoxifen (4-OHT; Sigma H7904) at concentrations ranging from 50–600 nM, prepared as 1000x stocks in ethanol for dose-optimization nuclear import studies. For nuclear export studies, cells were washed and treated with ethanol vehicle to enable reversal of 4-OHT-induced nuclear translocation. Real-time imaging of arCas9-^SPARK^-expressing cells was performed using a Nikon Eclipse E600 microscope.

### CRISPR editing assays

For validation of editing efficiency, cell lines co-expressing Cas9 were transduced or transfected with 6µg of pLenti CMV-EGFP –Blast^98^, and with LentiGuide-Puro sgRNAs targeting EGFP or ITGα6. The EGFP from LentiCRISPRv2EGFP-expressing cells was used in place of CMV-EGFP transfection. Cells were treated with 300 nM 4-OHT for 72 hours to induce editing and harvested 72 hours later to quantify EGFP or ITGα6 expression levels via flow cytometry. Results were analyzed with FlowJo v10 by gating for viable, singlet events, with editing efficiency calculated as the percentage decrease in EGFP or ITGα6-positive cells relative to non-targeting control sgRNA-transduced populations.

### Flow cytometry and fluorescence-activated cell sorting

Cells (1-2 x 10^6^) were detached using trypsin/EDTA (Gibco, 2500056) and stained in FITC-conjugated anti-integrinalpha6 (Abcam, ab30496) in FACS buffer (1% BSA, 0.5 mM EDTA, 25 mM HEPES in RPMI). Cells were interrogated using BD FACSCanto flow cytometer and quantification was performed with FlowJo software (BD Life Science). For enrichment of fluorescent-expressing cells (mCherry or EGFP), cells were facs sorted for fluorescent purity with a BD FACSAria II flow cytometer (BD Biosciences), expanded in 15-cm dishes and sorted anew for at least 3 consecutive rounds until they maintained greater than 80% purity. Cells with the highest MFI within the 80% pure cultures were isolated for stable cell line expansion in culture.

### arMAP algorithm for modeling arCas9 activity

The arMAP framework was used to predict editing efficiency solely based on inducible nuclear translocation dynamics. Cells expressing arCas9 and an EGFP reporter were exposed to 300 nM 4-OHT or vehicle. “OFF” baseline measurements were taken at 0 h; “ON” measurements were taken 72 h after induction. Editing efficiency (E) was quantified in parallel by EGFP fluorescence knockdown assessment using flow cytometry. Live singlets were gated, EGFP-positive frequency and median fluorescence intensity (MFI) were extracted, and *E* was defined as either %EGFP⁺ or EGFP-MFI normalized to vehicle controls, as indicated in each figure. Nuclear translocation (*T*) was quantified from arCas9^SPARK^ images acquired at matching time points using nuclear/cytoplasmic segmentation; *T* was expressed as the fraction (or mean N/C ratio) of cells exhibiting nuclear enrichment of arCas9^SPARK^’s EGFP signal. For arMAP model development, we fit a linear model relating translocation to editing:

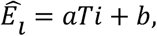

where 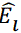 is the predicted editing efficiency at time point i, *Ti* is the corresponding translocation measurement, *a* is the slope, and *b* is an optional intercept capturing any baseline editing in the absence of translocation. Since arCas9 editing is negligible during the OFF-state, *b* is set to 0 and the model reduces to 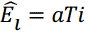. For experiments where only a 72h calibration time point is available, the slope is computed as

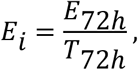

and then applied to earlier *Ti* to predict 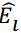 Linear regression was implemented in R using lm() with default settings; residuals were inspected for normality and homoscedasticity. Translocation time courses *Ti* were collected at multiple intervals after 4-OHT addition (e.g., 0, 1, 3, 6, 12, 24, 48, 72, and 180 min). arMAP predictions 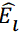 were generated for each *Ti* and compared to corresponding measured editing.

### Murine cSCC models and tumor cell line derivation

Transgenic *K14-CreER Notch1^L/+^ Pik3ca^E545K/+^ p53^fl/+^*and *K19-CreER Pik3ca^E545K/+^ p53^R172H/+^ Notch1^fl/fl^* mice were a gift from Matthew Ramsey. Autochthonous high-risk cSCC tumors were induced by treating mice with topical 4-hydroxytomoxyfen treatment (100 uL, 10mg/mL) on the shaved back during the telogen to 2^nd^ anagen transition (P25-30). In cases where tumor formation was not observed beyond 100 days, mice were treated once daily with 100 mg/kg tamoxifen (#T5648; Sigma-Aldrich) dissolved in sunflower seed oil (#S5007; Sigma-Aldrich) delivered by intraperitoneal injection for 5 days to activate Cre recombination as previously described^39^. To generate orthotopic transplants, autochthonous tumor tissues were minced and enzymatically digested with collagenase/hyaluronidase (STEMCELL Technologies), some cells were cultured *in vitro* to establish clonal tumor lines, while another portion was used for orthotopic graphs generated by subcutaneous injection of 1 x 10^6^ SCC cells with NIH3T3 cells 50/50 (v/v) PBS: Matrigel (Corning, 354234) into the flanks of NSG mice in a 50:50 mix of PBS and Matrigel as previously described^39^. Tumors were serially passaged twice orthotopically into NSG and C57BL/6 mice to enrich for metastatic phenotypes. Expression of epithelial markers was confirmed by immunofluorescence. Tumorigenicity was assessed via orthotopic re-implantation and histological staining for keratin pearls and hyperplasia. Tumors were measured daily, and tumor volumes were calculated using the following formula: tumor volume (mm^3^) = 4/3π × length/2 × width/2. For histology, Tumor tissues were fixed in formalin, embedded in paraffin, and sectioned at 4–5 µm thickness. Sections were stained with hematoxylin and eosin (H&E) for morphological analysis. Immunohistochemistry (IHC) was performed to assess markers like Ki-67 for proliferation and CD31 for vascularization. Immunofluorescence (IF) analysis was used to examine KRT14 and TP63 expression by the BWH histology core.

### Immunofluorescence and immunohistochemistry staining

Tumor sections were snap-frozen in HistoPrep (Fisher Scientific), and 5 um sections were cut. Sections and cultured cells were fixed in 3.7% Paraformaldehyde and permeabilized with 0.1% Triton X-100 followed by a block in 10% Normal Goat Serum, primary antibody staining and secondary antibody staining (AlexaFluor-488, AlexaFluor-568, Invitrogen). Nuclei were visualized by staining with Hoechst 33342 dye (Invitrogen). Samples were cover-slipped in Fluoromount mounting medium (Southern Biotech) and black and white images were obtained using a Nikon Eclipse E600 microscope and SPOT software. Pseudo colors were applied using Photoshop software (Adobe). The following antibodies were used for IF: K14 (Convave), p63 (4A4; Sigma-Aldrich; 1:20000), anti-CRISPR-Cas9 (7A9-3A3; Abcam; 1:200); for FACS: CD45 (biotin-conjugated, eBioscience), CD31 (biotin-conjugated, eBioscience), and CD49f (PE-conjugated; BD Biosciences). Processing of tumor samples was provided by the Dana-Farber/Harvard Cancer Center Specialized Histopathology Core Facility. Hematoxylin and eosin (H&E) stains were performed by the Harvard Medical Area Core Management System, Rodent Histopathology. Lung nodules were quantified by extracting lungs from tumor-bearing mice, inflating them with 70% ethanol, and preparing tissue for H&E sectioning. Nodules were counted as the number of keratinized tumor foci and nests per lung lobe. The total metastatic nodule burden per mouse was obtained by summing counts across all lobes.

### Clonal growth and spheroid formation assays

For cell colony density measurements, cells were plated at a density of 5 × 10³ cells into 10-cm culture dishes and allowed to grow for 10 days. At the endpoint, colonies were fixed in methanol at room temperature and stained with 10% Giemsa solution. For the scratch wound healing assay, 0.5× 10³ cells were cultured in 6-well plates to approximately 80% confluence in a humidified 5% CO₂ incubator at 37 °C. A single linear scratch was introduced using a sterile 200-μl pipette tip to simulate a wound. Detached cells were removed 24 h later by washing with PBS, and fresh medium was added. Wound closure was monitored every 12 h for 72 h using phase-contrast microscopy, and the percentage of wound closure was calculated relative to the initial wound width. 3D spheroid suspension assays were performed as previously described^47^. Briefly, 2 × 10⁴ cells were seeded into 96-well round-bottom plates pre-coated with a polymerized mixture of agarose and growth medium (1% w/v) and spheroid formation was assessed 24 h later. Soft agar assay was performed similarly to the previously published protocol^99^. Briefly, 2× 10⁴ cells were seeded for into 1.2% soft agar in individual wells of a 6-well plate. Cells were allowed to grow for 10 days then fixed in methanol and stained in crystal violet (0.0025%).

### Transwell migration and differentiation resistance assays

Cell lines’ invasive potential was measured as previously described^47^. Briefly, 2 × 10⁴ cells were seeded onto the upper chamber of a Matrigel matrix in a transwell dish (Corning 140620). Cells that invaded through the matrix and into the lower chamber were fixed in methanol and stained with 10% Giemsa solution and quantified under light microscopy. Keratinocytes resistance to differentiation signals was determined by maintaining 4 × 10⁴ cells per well in 6-well plates in KSFM supplemented with 1 mM calcium chloride and 5% FBS (DR medium)^100^. Cells were passaged every 3 days and replated at the same initial density into fresh DR medium for at least 12 passages.

### Construction of skin epigenome sgRNA library for pooled screening

A custom mouse skin-specific epigenome-wide sgRNA lentiviral library was designed to target 814 epigenetic modifiers and associated regulatory genes expressed in mouse skin. The library comprised a total of 2,672 sgRNAs, with four distinct sgRNAs per gene. In addition, 197 sgRNAs targeting essential genes (as defined by the Cancer Dependency Map^101^ were included as positive controls, and 365 non-targeting control (NTC) sgRNAs were included as negative controls. Library design, pooled oligonucleotide synthesis, library sequencing were performed by the Broad Institute’s Genetic Perturbation Platform (GPP) following established protocols (https://portals.broadinstitute.org/gpp/public/resources/protocols). The complete sgRNA sequences are provided in **Supplementary Table 1**. Pooled screening was conducted using previously published methods^102^. Briefly, 392-1 keratinocytes stably expressing either Lenti-arCas9-mCherry or LentiCRISPRv2 were transduced with the pooled lentiviral sgRNA library by spinfection (2 h, 931 × g) at a low multiplicity of infection (MOI 0.2) to ensure predominantly single-copy integration. Following transduction, cells were selected with puromycin until all uninfected NIC cells were eliminated. For RESTRICT-seq experiments, arCas9 activity was maintained in the OFF state during transduction and selection by omitting 4-OHT. Temporal activation of editing was achieved by treating cells with 300 nM 4-OHT in combination with 1 uM AZD4547 (Abcam) during defined therapeutic windows. At each passaging step, 4-OHT was withdrawn to restrict Cas9 activity to intended treatment intervals. Library representation was maintained at 500x coverage throughout the screen, with cell numbers adjusted accordingly at each passage. All screening experiments were conducted in phenol red–free, charcoal-stripped medium to eliminate exogenous hormone activity. Clonal growth assays were performed by plating 5 × 10³ cells in triplicate 150-mm dishes containing 30 ml medium. Cultures were fed every other day and serially passaged at low density every 3-4 days to determine cumulative population doublings (PD) over time. Cell number and viability at each passage were assessed using a Countess automated cell counter (Invitrogen, CA). PD was calculated using the formula: PD= (log N/N_0_)/log2, N = total cell # obtained at each passage and N_0_ = the number of cells plated at the start of the experiment.

### Data analysis for CRISPR screen

Cells were harvested at Days 3, 13, and 26 post-transduction, with >500× coverage maintained at all timepoints. Genomic DNA from screened cells was extracted using the DNA Blood Midi kit (Qiagen). Subsequent PCR and NGS sequencing were performed by the Broad GPP to determine sgRNA abundance. Temporal tracking of sgRNA abundance across long timescales was obtained from a sequenced pooled lentiviral library in which each sgRNA contained a sequence tag in the constant region, serving as a unique heritable barcode. Libraries were designed to avoid barcode collisions and synthesized by the Broad Institute GPP. Differential expression analysis was performed by MAGeck (version 0.5.9.5) and used to normalize the count table based on median normalization and fold changes in the conditions were calculated for genes and sgRNAs. Gene essentiality analysis was performed using MAGeCK Robust Rank Aggregation (RRA) to identify significantly enriched or depleted sgRNAs across experimental conditions. Raw sgRNA counts from PoolQ analysis served as input for MAGeCK RRA to assess the differential sgRNA abundance. To specifically evaluate drug-induced effects while accounting for baseline growth differences, we calculated normalized drug effects (normalized LFC) using the following approach (arCas9 groups: “Normalized drug effect = LFC(4-OHT AZD vs Vehicle [P8]) – LFC(4-OHT vs Vehicle [P8]) – LFC(Vehicle [P8] vs Vehicle [P0])”; wtCas9 groups: “Normalized drug effect = LFC(AZD vs Vehicle [P8]) – LFC(Vehicle [P8] vs Vehicle [P0])”. KEGG enrichment analyses were made using ShinyGO (version: v0.741; http://bioinformatics.sdstate.edu/go74/) with a p-value cutoff of 0.05. G.O. term data was processed through GO TermFinder (https://puma.princeton.edu/help/GO-TermFinder/GO_TermFinder_help.shtml). STARS negative binomial statistical framework (https://portals.broadinstitute.org/gpp/public/software/stars). For STARS analysis, PoolQ-normalized counts and corresponding.chip annotation files were used as inputs, with the analysis run in “Both” directions, a 10% sgRNA rank threshold, and default permutation-based FDR estimation to assign significance. This dual-analysis approach allowed robust hit calling by cross-validating results between the model-based MAGECK algorithm and the rank-based STARS scoring method, with high-confidence hits defined as those reproducibly enriched or depleted by both tools.

### Drug treatment, dose–response, and synergy assays

Drug synergy and dose–response assays were conducted using the following small-molecule inhibitors: PAK1 inhibitors AZ13705339 (Tocris, 6177) and NVS-PAK-1 (Tocris, 6132); FGFR inhibitor AZD4547 (Abcam, ab216311); KAT6A inhibitors WM-8014 (Tocris, 6693; ApexBio, A9779) and WM-1119 (MedChem Express, HY-102058); and CBP/P300 inhibitors SGC-CBP30 (Santa Cruz, 1613695-14-9), garcinol (Santa Cruz, 78824-30-3), GSK4027 (Sigma-Aldrich, SML2018), and its enantiomeric control GSK4028 (Sigma-Aldrich, SML2024). For IC₅₀ determination, cells were seeded in 96-well plates at 1×10^3^ cells per well and allowed to adhere overnight. The following day, cells were treated in triplicate with serial dilutions of each drug (ranging from 1000 µM to 0.01 µM) for 72 h in complete growth medium. Cell viability was quantified using the CellTiter Aqueous One Solution Cell Proliferation Assay (Promega, G3582), and absorbance was measured at 490 nm using a VICTOR³ plate reader (Perkin-Elmer, Waltham, MA, USA). Raw absorbance values were background-subtracted, drug concentrations were log₁₀-transformed (1000 µM = 3; 0.01 µM = –2), and response values were normalized to the percentage of vehicle-treated controls. Dose–response curves were fitted in GraphPad Prism using a nonlinear regression model with variable slope (four-parameter logistic). Best-fit values for LogIC₅₀, Hill slope, and IC₅₀ (µM) were obtained along with 95% confidence intervals via profile likelihood estimation. Model quality was assessed by R², sum of squares, and standard error of the regression (Sy.x). For synergy assays, cells were also treated in triplicate, but two different drugs were combined in an orthogonal matrix format in a 96-well plate, with each compound serially diluted (typically 0 µM to 50 µM) along either the X– or Y-axis, enabling evaluation of drug–drug interactions across a full range of concentration combinations.

### Statistical Analysis

All data were analyzed using GraphPad Prism v10.5.0 or R v4.5.1. For comparisons between two groups, unpaired two-tailed Student’s t-test or Mann–Whitney U test was used as appropriate. Multi-group comparisons were performed using one-way ANOVA followed by Tukey’s post-hoc test. Data are reported as mean ± standard deviation unless otherwise stated. Pearson’s correlation coefficient was used for linear associations. Statistical significance was defined as P < 0.05.

### Study approvals

All mice were housed and treated in accordance with protocols approved by the Brigham and Women’s Hospital Institutional Animal Care and Use Committee.

### Data access

The publicly available scRNA-seq data used in this study are accessible in the GEO database under accession codes GSE218170^83^. Sequencing data files generated in this study will be made available through in the Gene Expression Omnibus (GEO) database upon publication.

## ACKNOWLEDGEMENTS

We would like to acknowledge Selahattin Can Ozcan for his contributions to computational analyses, data interpretation, and editorial inputs. We would also like to thank the Broad Institute GPP, and Matt Ramsey (BWH) for assistance with technical troubleshooting, helpful comments, or consultation on this project. This work was supported by grants R00GM140262, K99GM140262; 1R21CA226099; 2P30CA013696-50; Columbia Data Science Institute Seeds Funds, and the Columbia HICCC-FAS Joint Pilot Award. Y.W was supported in part by 5T32AR007098.

## AUTHOR CONTRIBUTIONS

Y.W conceptualized, designed, and supervised the study in collaboration with the Broad Institute of MIT and Harvard. Y.W; D.A., Z.A., and A.N. performed the experiments. J.P., D.A., and Y.W., conducted the computational analyses. Y.W. and J.P., analyzed and interpreted the data. Y.W. wrote the manuscript. All authors read and approved the final manuscript.

## FIGURES

**SUPPLEMENTAL FIGURES**

**SUPPLEMENTAL TABLES**

**Supplemental Table 1. Epigenome sgRNA library**

**Supplemental Figure 1.**
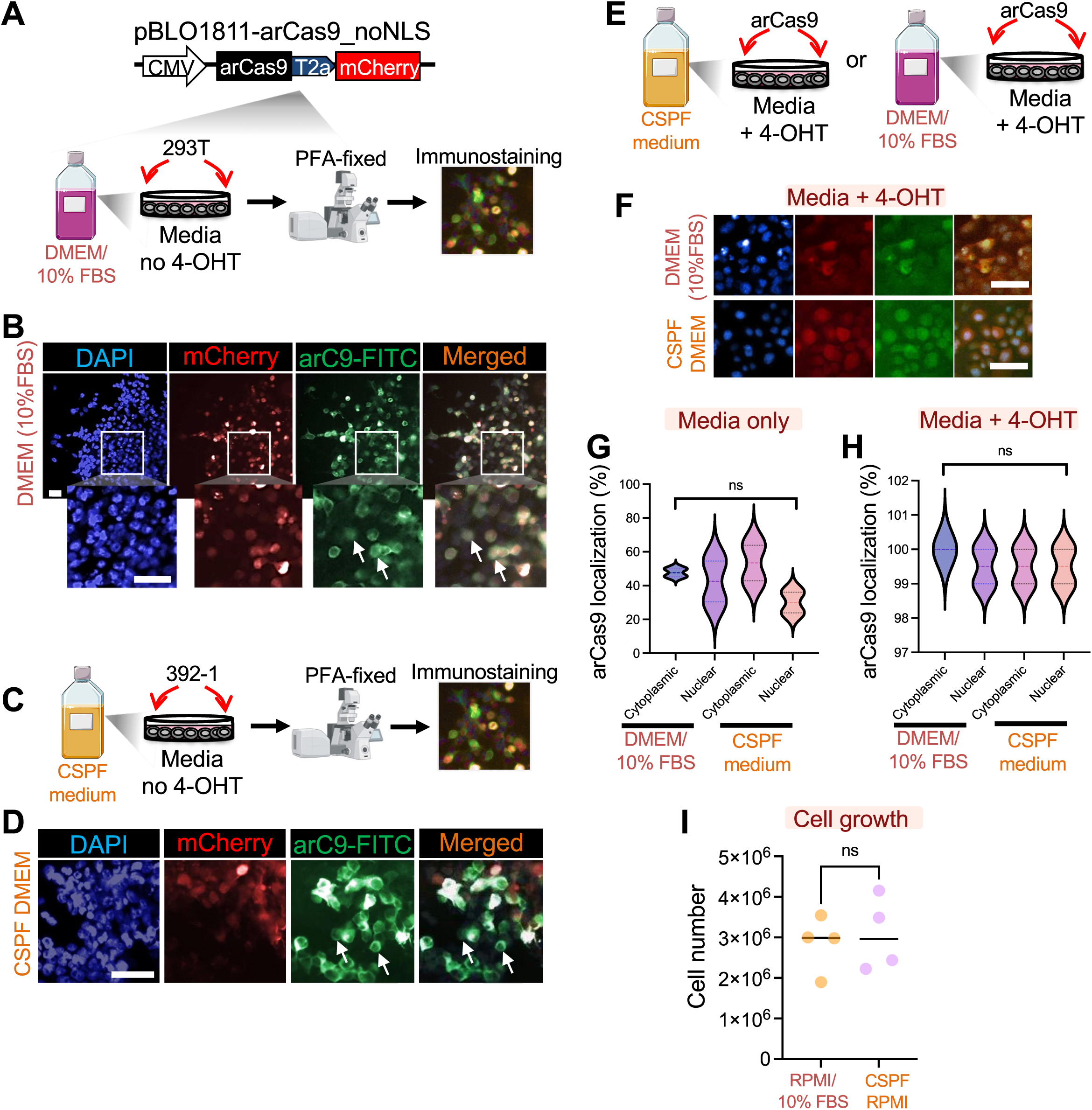
Charcoal-stripped medium attenuates arCas9 background nuclear translocation. (A) Schematic of pBLO1811-arCas9mCherry-noNLS vector transfection into 293T cells cultured in DMEM with 10% FBS. The intracellular location of the arCas9 protein is subsequently assessed by immunofluorescence staining to visualize the distribution of arCas9 in the absence of 4-OHT. (B) Representative images of 293T cells immunostained for DAPI (nuclear stain), mCherry (arCas9-t2a expression), and Cas9 to confirm arCas9 presence. The boxed area highlights cell colonies with increased arCas9 nuclear protein levels. White arrows point to individual cells showing elevated background nuclear translocation. The white box designates zoom-in area shown in the bottom panel. *n* = 2 technical replicates per experiment in two representative cell lines. Scale bars, 50 µm (C) Schematic depicting the expected intracellular location of arCas9 in mammalian cells cultured in charcoal-stripped, phenol-red-free medium designed to minimize estrogenic effects that could lead to unspecific nuclear translocation. (D) Representative images of 293T cells immunostained for DAPI, mCherry (arCas9-t2a expression), and Cas9. White arrows indicate individual cells with high cytoplasmic arCas9 protein. *n*= 2 technical replicates per experiment in one representative cell line. Scale bars, 50 µm. (E) Schematic of the arCas9 intracellular location assay in 4-OHT-treated cells cultured in either charcoal-stripped phenol-red-free medium (CSPF) or unstripped DMEM. (F) Images of 4-OHT-treated 293T cells immunostained for DAPI, mCherry (arCas9-t2a expression), and Cas9. *n* = 2 technical replicates per experiment in two representative cell lines. Scale bars, 50 µm. (G-H) Quantification of cells displaying cytoplasmic and nuclear arCas9 compartmentalization before or after 4-OHT treatment in DMEM with 10% FBS or CSPF-conditioned medium. *n* = 2 technical replicates per experiment in two representative cell lines. Statistical significance was determined using one-way ANOVA plus Tukey’s MC test (ns: not significant). (I) Quantification of 392-1 cell growth in standard RPMI supplemented with 10% FBS versus CSPF-conditioned RPMI medium

**Supplemental Figure 2.**
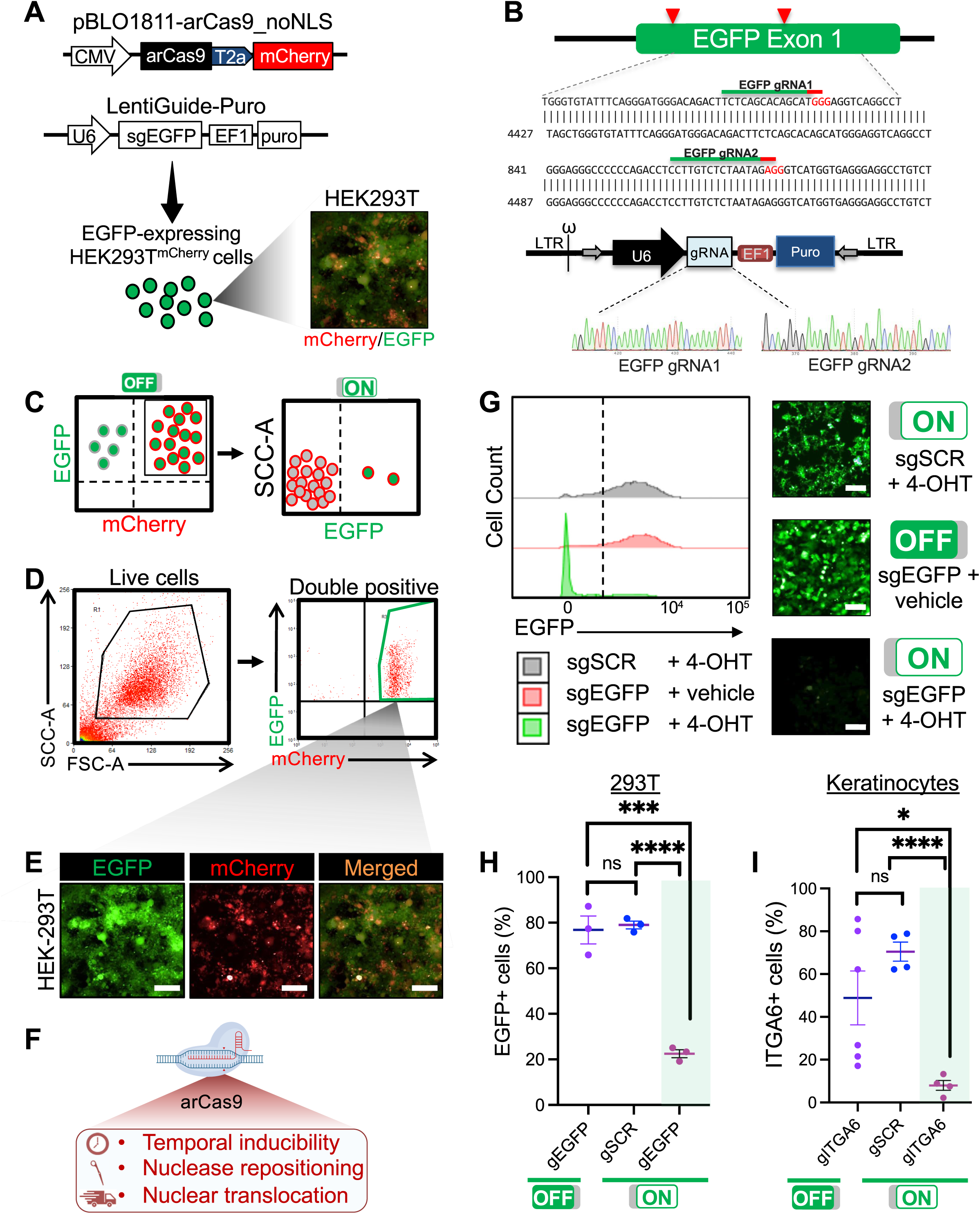
Validation of arCas9 inducibility and catalytic activation for gene silencing in mammalian Cells. (A) Schematic representation of the Lenti-arCas9 and LentiGuide-Puro plasmids used in the EGFP gene disruption assay. The Lenti-arCas9 construct enables expression of the allosterically-regulated Cas9 (arCas9), which is catalytically inactive in the absence of 4-OHT for targeted gene repression. (B) The LentiGuide-Puro plasmid delivers two separate gRNAs, both designed to target exon 1 of the EGFP gene. gRNA targeting sequences are confirmed by Sanger sequencing, with sequences highlighted in red indicate the protospacer adjacent motif (PAM) necessary for Cas9 binding and cleavage. (C) Schematic depicting the EGFP gene repression assay, where arCas9 is catalytically activated with 4-OHT leading to a conformational change that enables DNA cleavage as measured by flow cytometry. (D) Fluorescence-activated cell sorting (FACS) purification of EGFP-positive, arCas9-mCherry co-expressing HEK-293T cells co-transfected with the Lenti-arCas9-mCherry and the pLL3.7-EGFP plasmids. (E) Representative live immunofluorescent images of HEK-293T arCas9-mCherry/EGFP reporter lines (293T-mCherry) purified by FACS isolation. Scale bars, 50 µm. (F) Three distinct mechanisms of regulation of arCas9 catalytic activation controlled by 4-OHT ligand binding: 1) Temporal inducibility (activation only occurs in the presence of 4-OHT); 2) Repositioning of the HNH and RuvC nuclease domains for DNA cleavage; 3) Nuclear translocation of arCas9 (where it can interact with genomic DNA). (G) Representative flow cytometric and immunofluorescent assessment of HEK-293T arCas9-mCherry/pLL3.7-EGFP reporter lines transfected with scramble gRNA (sgSCR) or an EGFP targeting gRNA (sgEGFP) then exposed to 300 nM 4-OHT or vehicle for 72h. *n* = 3 technical replicates from one representative cell line. Scale bars, 50 µm. (H) Flow cytometric measurements of 293T-mCherry-EGFP and (I) N/TERT keratinocytes edited by sgEGFP, sgSCR, or sgITA6, respectively, for 72 h using catalytically active (ON) or inactive (OFF) arCas9 in standard DMEM/10% FBS media supplemented with 300 nM 4-OHT. Data are representative of two independent experiments with at least *n* = 3 technical replicates per sgRNA. Bar graphs represent the mean; error bars represent standard deviation (s.d). Significances were determined using a two-sided Student’s unpaired t-test (*: p < 0.05, ***: p < 0.001, ****: p < 0.0001).

**Supplemental Figure 3.**
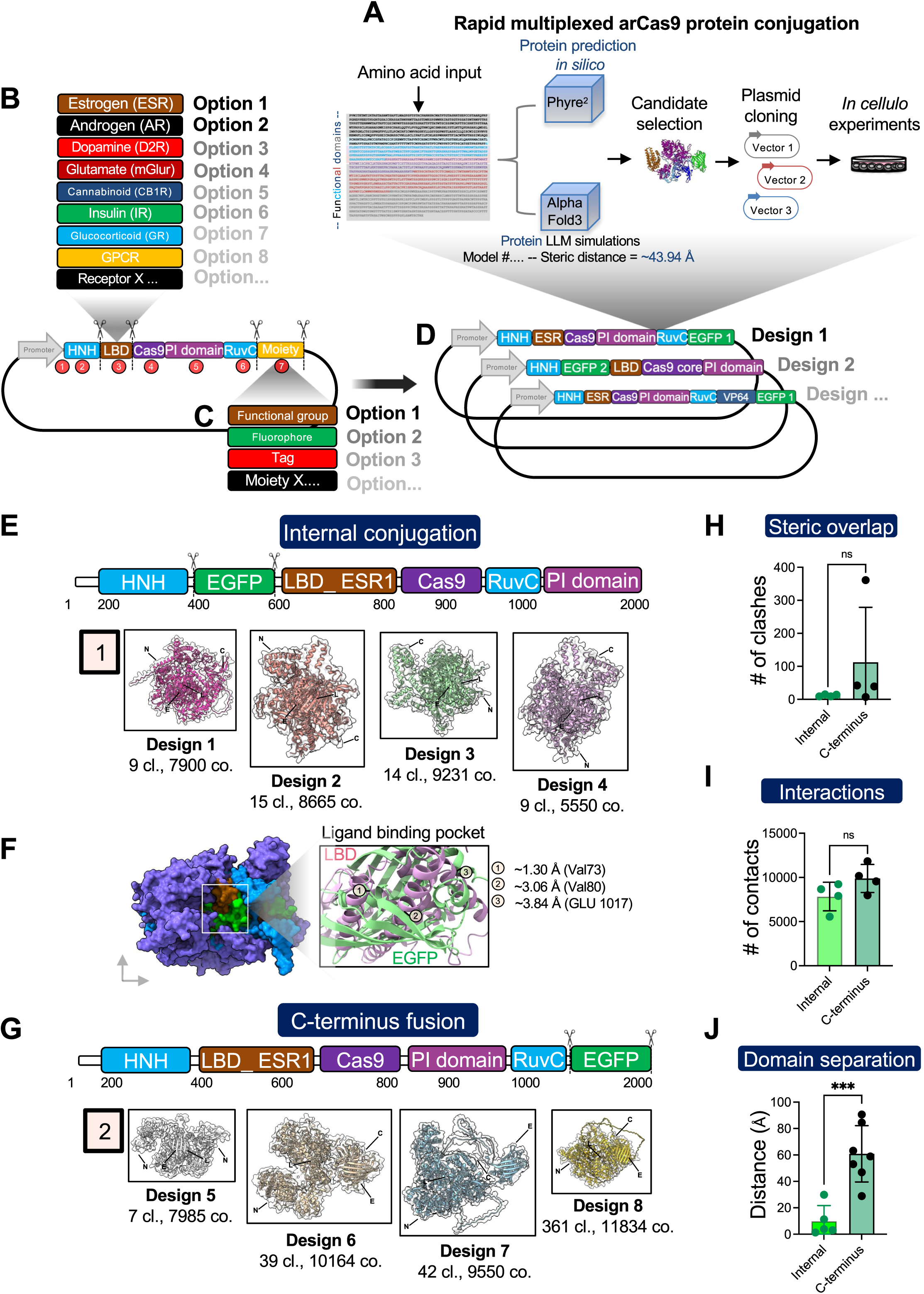
*In silico* platform for engineering multiplexed functional domain insertions in arCas9. (A) Schematic overview of the pipeline used for engineering and testing additional protein insertions into arCas9. This modular design facilitates the integration of various allosterically regulated receptors and functional groups, enhancing the versatility of arCas9 for tailored molecular applications. (B) Assorted ligand binding domains with allosteric functions can be substituted for the ligand-binding domain (LBD) at vector position 3, allowing for customized tuning of arCas9’s regulatory mechanisms. Scissors indicate highly modular insertion sites designed to accept various functional domain conjugations with minimal disruption to arCas9 allosteric functions. (C) Additional tailored functional groups can be seamlessly integrated by insertions at the C-terminus on vector position 7. (D) Plasmid library stack rationally engineered to include EGFP along with other functional groups for real-time fluorescent tracking of protein allosteric activity, providing visual means to monitor the dynamic behavior of arCas9. (E) Mapping arCas9-EGFP protein fusion engineered by internal conjugation of EGFP near the LBD at vector position 3. Amino acid positions for the HNH and RuvC endonuclease domains are designated in blue, EGFP in green, estrogen receptor 1 (ESR1) LBD in sepia, Cas9 core protein in purple, and the protospacer adjacent motif interacting (PI) domain in violet. (1) Protein simulation showing the N-terminus (N), C-terminus (C), EGFP (E), and ligand-binding domain (L). The number of protein clashes (cl) and contacts (co) are indicated below each structure to assess potential steric hindrance or beneficial interactions. (F) 3D protein structure of the ligand binding pocket (LBD) of an internally-conjugated arCas9-EGFP variant, displaying at least 3 obstructive clashes between LBD and EGFP, as visualized in ChimeraX. (G) Mapping arCas9-EGFP protein fusion engineered by C-terminal conjugation of EGFP at vector position 7. (2) Simulations of arCas9-EGFP C-terminal protein fusions. Protein clashes (cl) and contacts (co) are indicated below each structure. (H) Quantification of steric overlap as measured by the number of unfavorable clashes between atoms of the arCas9-EGFP protein engineered by internal or C-terminus fusion of EGFP. (I) Quantification of residue interactions as measured by the number of contacts between atoms of the arCas9-EGFP protein engineered by internal or C-terminus fusion of EGFP. (J) Quantification of domain separation as measured by total distance between the LBD and EGFP functional groups in the arCas9-EGFP fusion protein engineered by internal or C-terminus fusion. Bar graphs represent the mean ± SD of *n* = 4-7 replicates. Significances were determined using a two-sided Student’s unpaired t-test (*: p < 0.05, **: p < 0.01, ***: p < 0.001, ****: p < 0.0001).

**Supplemental Figure 4.**
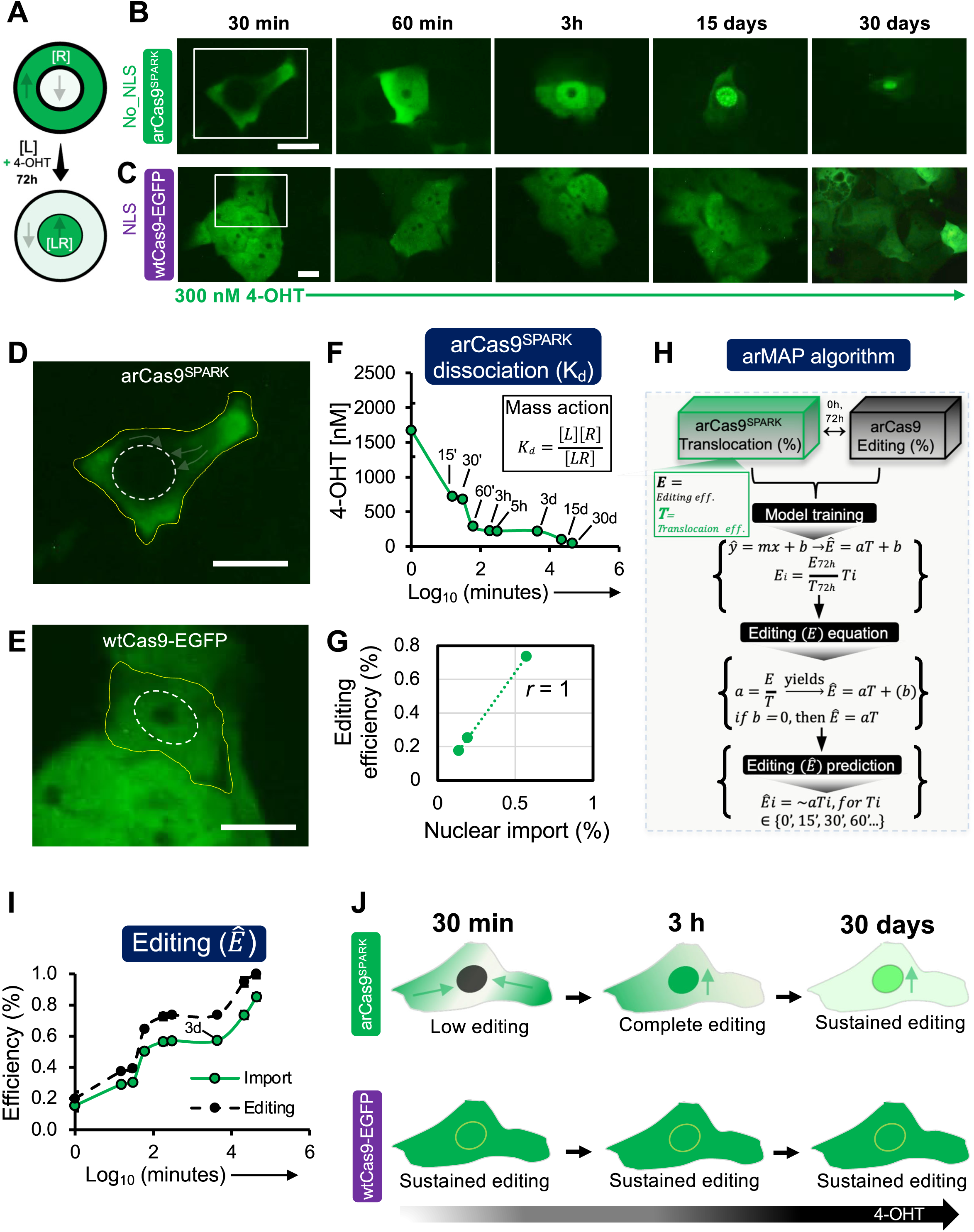
arCas9SPARK calibrates arCas9’s sustained long-term nuclear compartmentalization and editing. (A) Theoretical overview of the law of mass action variables generated by ligand [L] concentration, unbound arCas9^SPARK^ concentration [R], and the concentration of the bound arCas9^SPARK^–ligand complex [LR]. (B) Representative images of a time series analysis in Lenti-arCas9^SPARK^ –expressing SCC-13 cells treated with 300 nM 4-OHT. (D)The white box designates areas zoomed in the bottom panels. Scale bar, 10μm. *n* = 3 technical replicates from one representative cell line. (C) Representative images of SCC-13 cells expressing LentiCas9CRISPRv2-EGFP treated with 300 nM 4-OHT for comparison with arCas9^SPARK^. The white box designates areas zoomed in the bottom panels (E). Scale bar, 10μm. *n* = 3 technical replicates from one representative cell line. (D) Representative image of a SCC-13 cells expressing arCas9^SPARK^ treated with 300 nM 4-OHT for 30 minutes capturing the initial phase of nuclear translocation. The white outline depicts the nucleus, and the yellow outline demarcates the cell membrane. Arrows indicate movement patterns of the EGFP signal towards the nucleus to visually compare the dynamics of EGFP localization between the two Cas9 systems. Scale bar, 10μm. (E) Representative image of a SCC-13 cells expressing Lenti-Cas9CRISPR-EGFP treated with 300 nM 4-OHT for 30 minutes. The white outline depicts the nucleus, and the yellow outline demarcates the cell membrane. Scale bar, 10μm. (F) Graph of arCas9^SPARK^ dissociation constant over 30 continuous days (4.32×10^4^ minutes) of exposure to 300 nM 4-OHT, demonstrating the stability of ligand binding over an extended period. (G) Correlation between editing efficiency and nuclear import under three conditions: (1) arCas9-OFF with a gene-targeting gRNA, (2) arCas9-ON with a gene-targeting gRNA. Data represent the mean of three technical replicates from a representative cell line. (H) arCas9 Modeling for Activity Prediction (arMAP) algorithm which computes predictions of arCas9 editing efficiency based solely on correlated arCas9^SPARK^ translocation trends. (I) Predicted arCas9 editing efficiencies (*Ê*) derived from corresponding arCas9^SPARK^ translocation ($) timepoints. Bar graphs represent the mean ± SD of *n* = 3 technical replicates from one representative cell line. Statistical significance was determined using a two-sided Student’s unpaired t-test. (J) Graphical representation of editing and nuclear translocation dynamics in wtCas9 and arCas9 systems. Unlike conventional wtCas9 systems, arCas9 enables precise, ligand-dependent control of nuclear import and gene editing, minimizing background activity prior to induction.

**Supplemental Figure 5.**
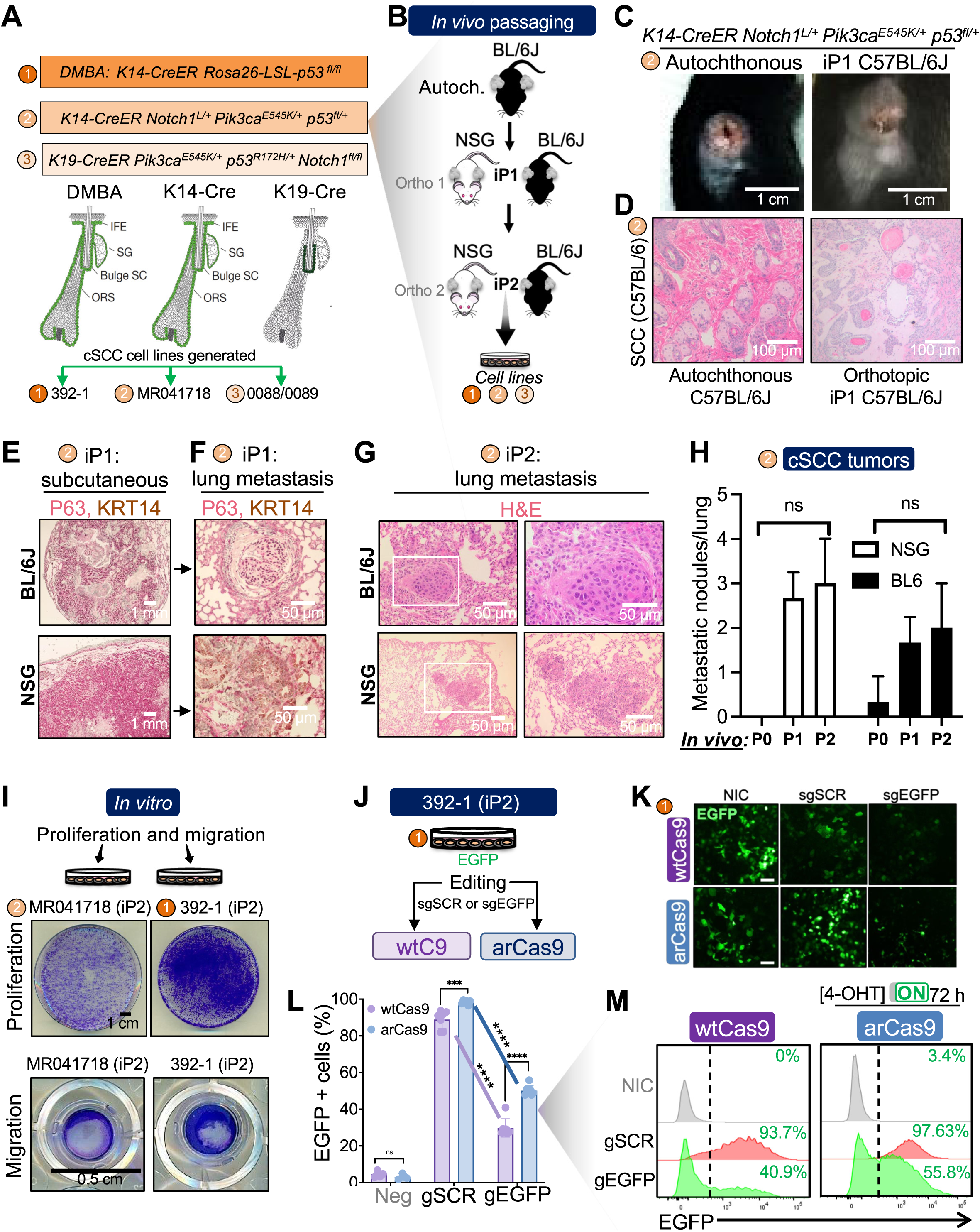
Orthotopic serial passaging of autochthonous SCC tumors recapitulates high-risk cSCC. (A) Schematic depicting three mouse models used to generate autochthonous tumors from distinct skin epithelial compartments: (1) DMBA-induced *K14-CreER, Rosa26-LSL-P53^fl/fl^*, modeling skin cancer with p53 loss. (2) *K14-CreER, Notch1^L/+^ Pik3ca^E545K/+^ p53^fl/+^*, combining activating mutations in *Notch1* and *PIK3CA* with conditional p53 knockout. (3) *K19-CreER*, *Pik3ca^E545K/+^ p53^R172H/+^ Notch1^fl/fl^*, targeting a different epithelial compartment with a PIK3CA mutation, a p53 hotspot mutation, and Notch1 activation. The respective cell lines obtained from each model are also indicated. (B) Illustrative depiction of *in vivo* tumor passaging assay used to establish HR-cSCC cell lines. (C) Representative images of autochthonous and first *in vivo* passaged (iP1) tumors from *K14-CreER, Notch1^L/+^ Pik3ca^E545K/+^ p53^fl/+^* mice with (D) corresponding H&E histology to show tumor morphology and cellular architecture. *n* = 3 biological replicates. Scale bars are indicated by white bars. (E) Immunohistochemistry (IHC) analysis of keratin 14 (KRT14) and tumor protein 63 (TP63) expression in tumors derived from *K14-CreER, Notch1^L/+^ Pik3ca^E545K/+^ p53^fl/+^*mice after one passage in C57BL/6 or NSG mice to confirm retention of skin basal cell markers post-passage. (F) KRT14 and TP63 IHC analysis of corresponding lung samples to detect metastatic spread. Scale is depicted with white bars. (G) Hematoxylin & eosin (H&E) staining of lung samples harvested from C57BL/6 or NSG mice inoculated with tumors from *K14-CreER, Notch1^L/+^ Pik3ca^E545K/+^ p53^fl/+^* mice at the second *in vivo* passage (iP2). The white box designates the area zoomed in the right panel for detailed examination. (H) Quantification of lung metastatic nodules in NSG and C57BL/6 mice inoculated with *K14-CreER, Notch1^L/+^ Pik3ca^E545K/+^ p53^fl/+^* across multiple *in vivo* passages. *n* = 3 biological replicates. (I) Representative images of cell colony density and transwell migration examined by Giemsa staining in DMBA tumor-derived 392-1 cell line and *K14-CreER, Notch1^L/+^ Pik3ca^E545K/+^ p53^fl/+^* tumor-derived MR041718 cell line. (J) Schematic of EGFP disruption experiments in 392-1-EGFP cells stably expressing either wild-type Cas9 (wtCas9) or arCas9. (K) Immunofluorescent images showing EGFP disruption in 392-1 cells at 72 hours post-transfection with either wtCas9 or 300 nM 4-OHT-treated arCas9. *n* = 3 technical replicates from one representative cell line. Scale bar, 50μm. (L) Flow cytometric quantifications and (M) representative histogram graphs of EGFP repression in 392-1 cells stably expressing wtCas9 or arCas9, treated with either scramble gRNA or EGFP-targeting gRNA, relative to non EGFP-expressing negative control for 72 hours in standard RPMI/10% FBS media supplemented with 300 nM 4-OHT. Bar graphs represent the mean ± SD of *n* = 3-7 technical replicates from three separate experiments. Statistical significance was determined using a two-sided Student’s unpaired t-test for direct comparisons or one-way ANOVA followed by Tukey’s Multiple Comparison (MC) test for multiple group comparisons (*: p < 0.001, ****: p < 0.0001).

**Supplemental Figure 6.**
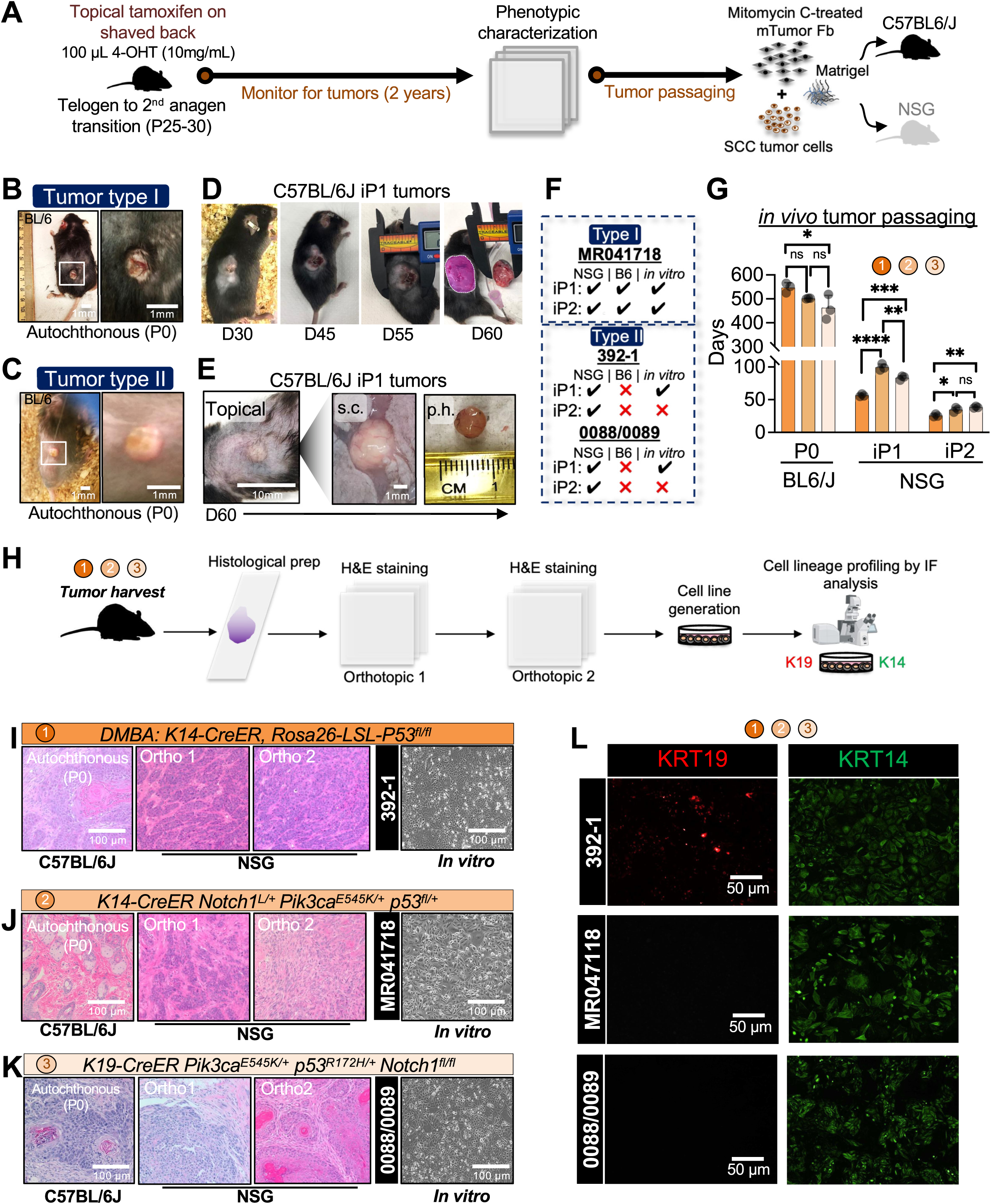
Characterizing high-risk cSCC mouse models. (A) Experimental outline of the protocol used for generating skin-specific autochthonous tumors and the subsequent *in vivo* serial transplantations in mice. (B-C) Representative images of autochthonous tumor types I and II. The white box designates an area zoomed in on the right panel for detailed view. Scale is indicated by white bars. (D) Representative dorsal view of a mouse displaying a prominent exophytic type I tumor after one round of *in vivo* passage (iP1), maintained for over 60 days. Scale bar, 100 μm. (E) Representative dorsal view of exophytic type II tumors after one round of *in vivo* passage, maintained for over 60 days. Tumors are shown topically, subcutaneously (s.c.), and post-harvest (p.h.). Scale bar, 100 mm or as indicated. (F) Mouse tumor cell line growth outcomes for type I (endophytic) and type II (exophytic) tumors in NSG or C57BL/6 mice after two rounds of *in vivo* passages. Checkmarks indicate successful *in vivo* grafting or *in vitro* plating, summarizing the ability of different tumor types to establish as cell lines. (G) Tumorigenicity analysis measuring days to tumor formation after cell inoculation of tumor cells from: (1) DMBA-induced *K14-CreER, Rosa26-LSL-P53^fl/fl^*. (2) *K14-CreER, Notch1^L/+^ Pik3ca^E545K/+^ p53^fl/+^*. (3) *K19-CreER, Pik3ca^E545K/+^ p53^R172H/+^ Notch1^fl/fl^* tumors during successive *in vivo* passaging. *n* = 3 biological replicates. Bar graphs represent the mean ± SD of *n* = 3 biological replicates from three separate experiments. Statistical significance was determined using one-way ANOVA followed by Tukey’s Multiple Comparison (MC) test (*: p < 0.05, **: p < 0.01, ***: p < 0.001, ****: p < 0.0001). (H) Schematic illustration of the histological and cell lineage analysis conducted on cSCC mouse tumors, outlining the methodology used for characterizing tumor histology and cellular origin. (I-K) Histological analysis of tumors from (1) DMBA-induced *K14-CreER, Rosa26-LSL-P53^fl/fl^*. (2) *K14-CreER, Notch1^L/+^ Pik3ca^E545K/+^ p53^fl/+^*. (3) *K19-CreER, Pik3ca^E545K/+^ p53^R172H/+^ Notch1^f/fl^*after two successive orthotopic passages. (Right) Representative phase contrast images of corresponding *in vitro* cell lines generated and maintained in keratinocyte growth media. Scale is indicated by white bars. (L) Immunofluorescence analysis (IF) of keratin 14 (KRT14) and keratin 19 (KRT19) expression in cell lines established from mouse-derived tumors at passage 3 *in vitro*.

**Supplemental Figure 7.**
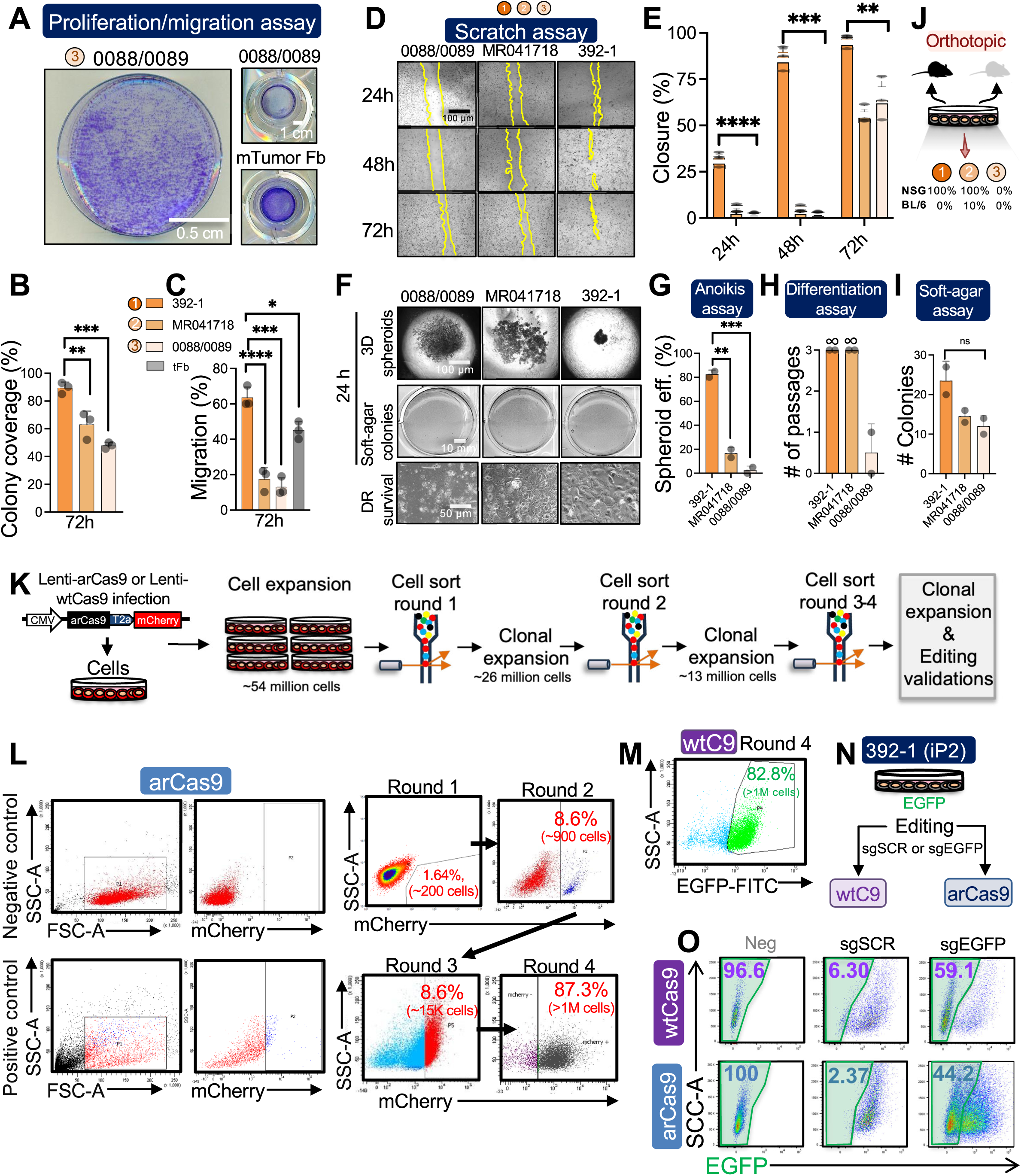
Characterization of mouse cSCC tumor cell lines and generation stable wtCas9 and arCas9 expressing derivatives. (A) Representative images of cell colony density and transwell migration assessed by Giemsa staining in 0088/0089 cell line derived from *K19-CreER, Pik3ca^E545K/+^ p53^R172H/+^ Notch1^fl/fl^* cSCC tumors and mouse tumor fibroblast controls. Scale depicted with white bars. (B) Quantification of colony coverage and (C) transwell migration rates across various mouse tumor cell lines at 72 hours post-incubation. (D) Representative images and (E) quantification of migration zones for each mouse tumor cell line captured over a 72-hour period. (F) Representative images and (G-I) quantifications of self-assembled suspension spheroid formation, soft agar colony formation, and cell survival in differentiation-inducing media across mouse cSCC tumor lines. *n* = 2 replicates from each cell line. (J) Rate of successful orthotopic tumor grafting generated from *in vitro* cultures of mouse cSCC lines derived from (1) DMBA-induced *K14-CreER, Rosa26-LSL-P53^fl/fl^*; (2) *K14-CreER, Notch1^L/+^ Pik3ca^E545K/+^ p53^fl/+^*; (3) *K19-CreER, Pik3ca^E545K/+^ p53^R172H/+^ Notch1^fl/fl^* tumors. (K) Workflow of stable cell line generation and cell line purification for Lenti-arCas9-mCherry and LentiCas9CRISPRv2-EGFP (wtCas9)-expressing cells. (L-M) Representative flow cytometric gating strategy employed for isolating Lenti-arCas9-mCherry and wtCas9-EGFP-expressing cells over 4 rounds of serial purification and expansion. (N) Schematic of EGFP disruption experiments in EGFP-expressing mouse tumor cells stably expressing wtCas9 or arCas9. (O) Representative flow cytometry plots showing EGFP disruption after 72 hours in 392-1 cells transduce with either scramble gRNA or EGFP-targeting gRNA. Comparisons include wtCas9-expressing cells and arCas9-expressing cells treated with 300 nM 4-OHT. Significances were determined using a one-way ANOVA plus Tukey’s MC test (*: p < 0.05, **: p < 0.01, ***: p < 0.001, ****: p < 0.0001).

**Supplemental Figure 8.**
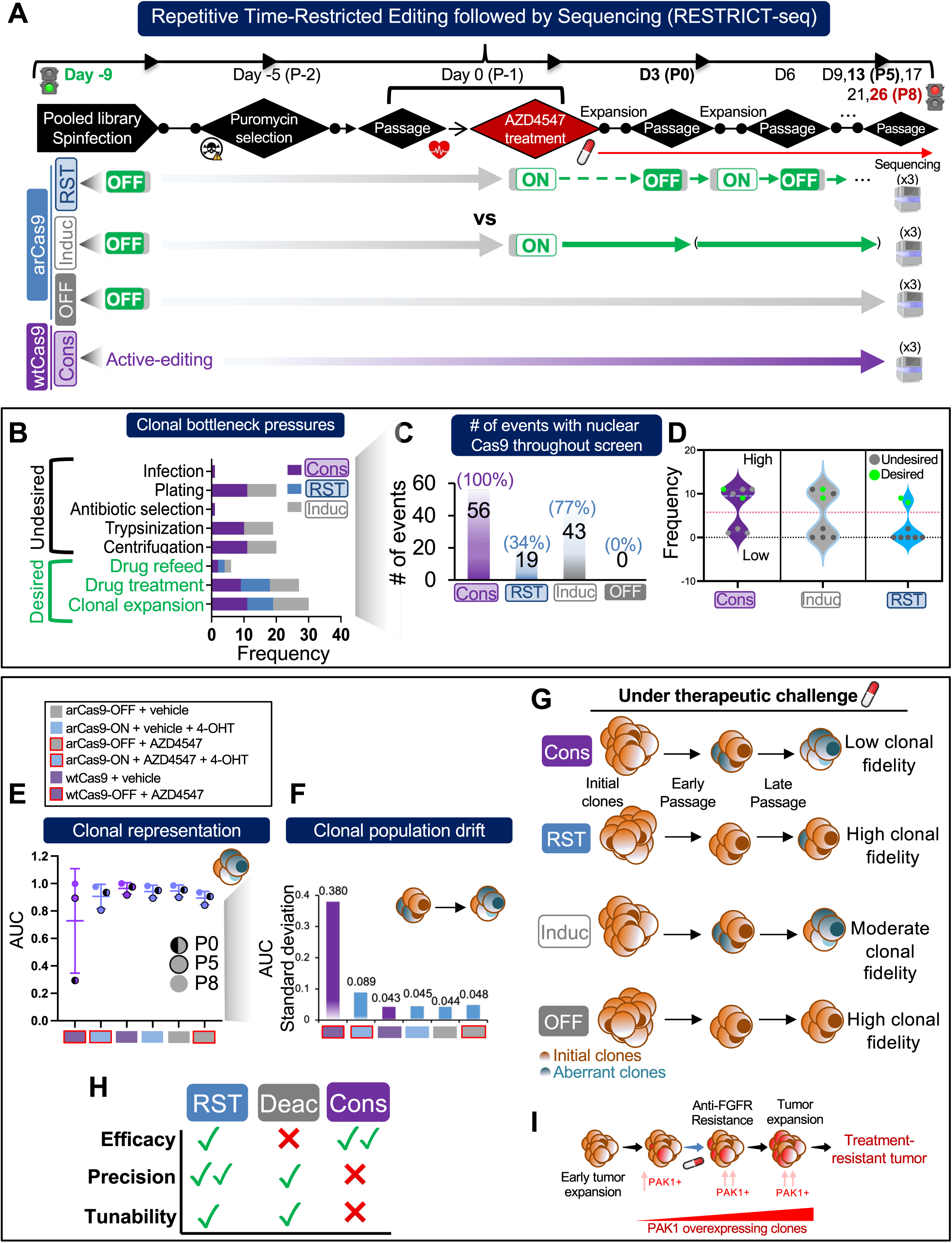
RESTRICT-seq eliminates undesired disruptions in pooled screening and mitigates population clonal drift. (A) Experimental design of the Repetitive Time-Restricted Editing followed by Sequencing (RESTRICT-seq) protocol, compared to traditional pooled CRISPR screening methods e.g. Constitutively-active wtCas9 screens (Cons): where Cas9 is active throughout the screen; Inducible arCas9 screens (Ind): where Cas9 activity is controlled by the presence of an inducer (e.g., 4-OHT) early in the screen; Deactivated arCas9 screen (OFF): where Cas9 remains catalytically inactive throughout the screen, serving as a control for background effects. Bolded days indicate time points when sequencing was performed. (B) Categorical quantification of desired and undesired clonal selection events accumulated during a 35-day long pooled dropout screen across all pooled screening conditions. (C) Quantification of the total number of clonal selection events during which Cas9 was catalytically active. (D) Quantification of undesired and desired clonal disruptions under catalytically active Cas9s: constitutive wtCas9 screens (Cons), inducible arCas9 screens (Induc), and RESTRICT-seq (RST) CRISPR pooled screens. Events above the red line represent high-frequency disruptions with the greatest impact on population clonal drift. (E) Area under the curve (AUC) analysis of cumulative gRNA clonal abundance across all pooled screening conditions, encompassing three passage time points. (F) Standard deviation measurements of AUC depicting clonal drift across all pooled screening conditions. (G) Schematic representation of clonal selection biases generated by various pooled CRISPR dropout screen protocols. (H) Schematic summary of the overall impact of each screening method on maintaining true clonal representation. (I) Theoretical overview of early SCC tumor clonal selection depicts PAK1 overexpression as an early adaptation for anti-FGFR therapy resistance in treatment-naïve tumors.

**Supplemental Figure 9.**
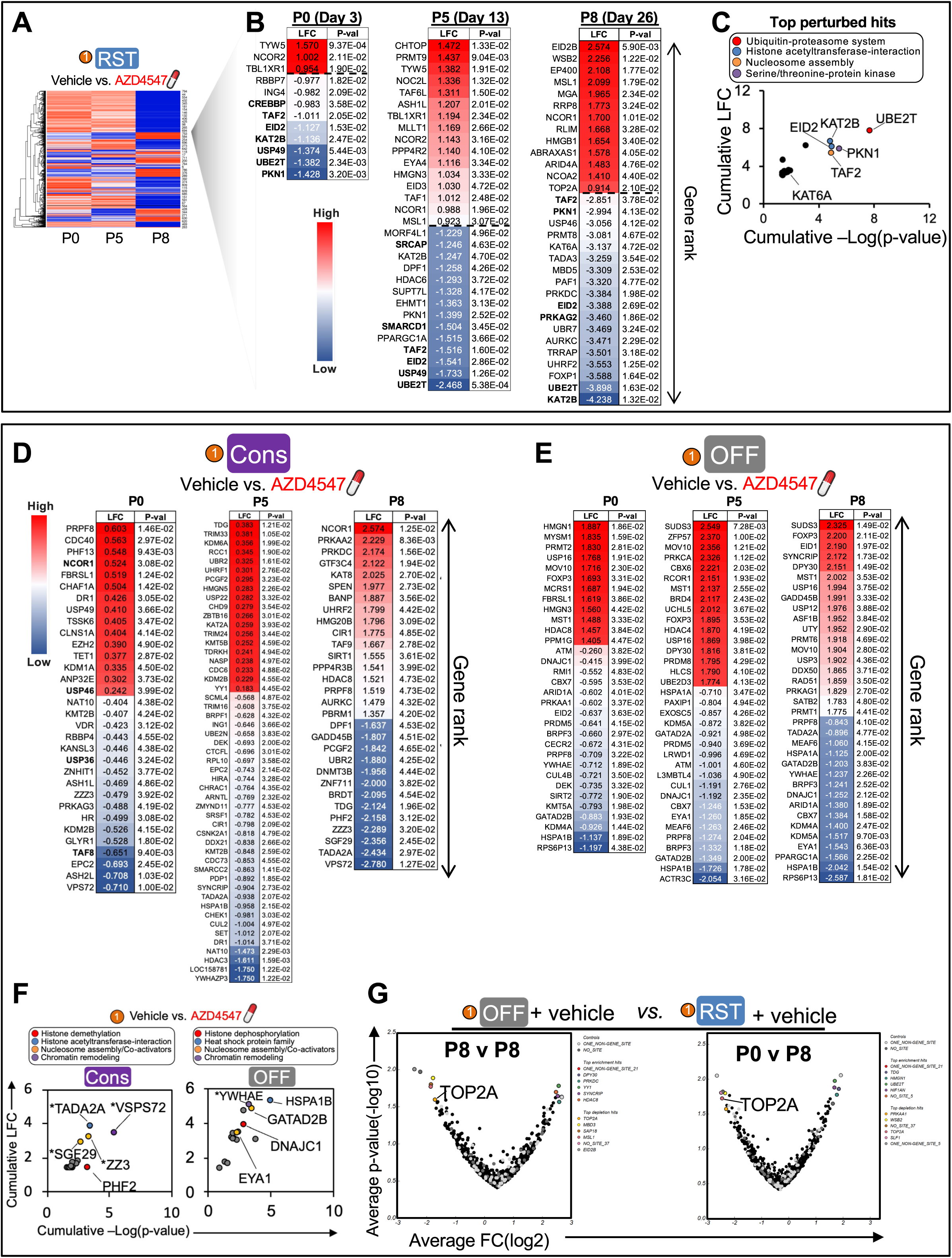
Constitutive wtCas9 screens shows biased overrepresentation of essential genes over drug-synergistic hits. (A-B) STARS hypergeometric ranking of significantly enriched or depleted target genes in the RESTRICT-seq screen under AZD4547 versus vehicle at P0, P5, and P8. *n* = 3 technical replicates; sgRNA ranks were analyzed based on significant average –log₁₀(p) value. (C) Quantification of cumulative LFC versus the negative log p-value for significantly depleted target genes in the AZD4547-treated RESTRICT-seq screen. Colors differentiate associated gene programs. (D) Ranking of significantly perturbed target genes with a p-value ≤ 0.05 and a log fold change (LFC) of ± 0.9 in the AZD4547-treated Cons screen at all time points (P0, P5, P8). (E) Ranking of perturbed target genes with a p-value ≤ 0.05 and a log fold change (LFC) of ± 0.9 in the AZD4547-treated arCas9-OFF screen at all time points (P0, P5, P8). (F) Quantification of cumulative LFC versus the negative log p-value for significantly depleted target genes in AZD4547-treated Cons and OFF screens. Colors differentiate associated gene programs. Asterix demarcate essential genes. (G) Volcano plots comparing significantly perturbed genes between vehicle-treated groups from RESTRICT-seq and arCas9-OFF screens, between P0 to P5 and P0 to P8.

**Supplemental Figure 10.**
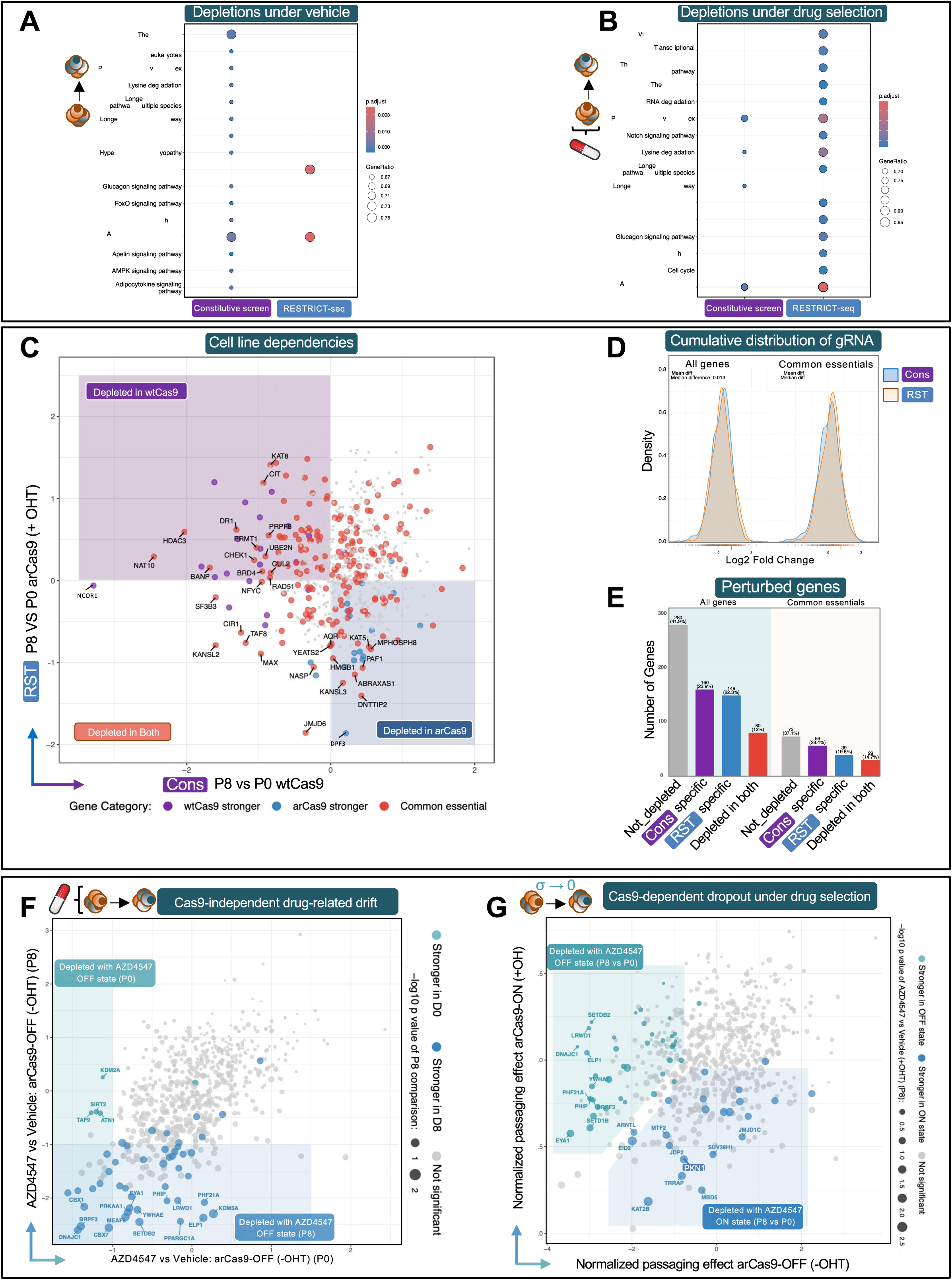
RESTRICT-seq enhances gene dropouts related to Cas9-dependence and drug synchrony. (A) KEGG pathway enrichment analysis of genes significantly depleted in vehicle-treated constitutive wtCas9 and arCas9-ON screens at passage 8 (P8) relative to passage (0) P0. (B) KEGG pathway enrichment analysis of genes significantly depleted in AZD4547-treated constitutive wtCas9 and RESTRICT-seq screens at passage 8 (P8) relative to vehicle-treated controls. (C) wtCas9– and arCas9-specific depletions at P8 compared to P0 in vehicle-treated cells. Top 30 depletions are labelled on plot. (D) LogFC distribution of gRNAs in constitutive wtCas9 and RESTRICT-seq screen under vehicle treatment. (E) Distribution of gene depletions in vehicle-treated constitutive wtCas9 and RESTRICT-seq screens at passage 8 (P8) relative to passage 0 (P0). (F) Comparative Robust Rank Aggregation (RRA) analysis of depleted genes from AZD4547-treated arCas9-OFF CRISPR screens at P0 and P8, relative to vehicle-treated controls. (G) RRA analysis of depleted genes from AZD4547-treated RESTRICT-seq vs arCas9-OFF CRISPR screens at P8 vs baseline (P0), relative to vehicle-treated controls.

**Supplemental Figure 11.**
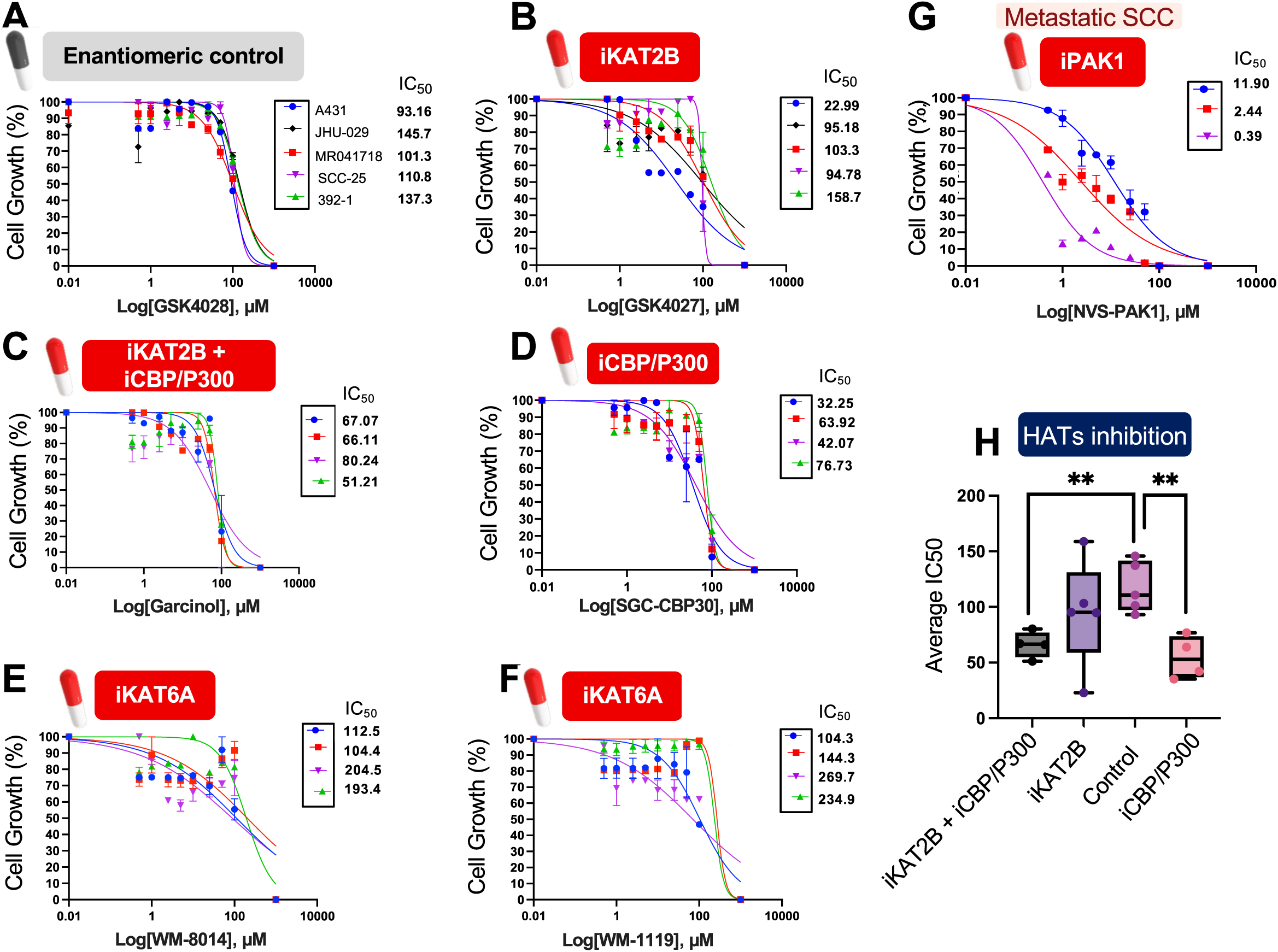
Inhibition of Histone Acetyl Transferases Alone Displays Modest Reduction in Tumor Cell Survival. (A) 10-point dose-response small molecule inhibition screen and IC50 determinations with the enantiomeric inhibitor control (GSK-4028), providing a baseline for non-specific effects in human and mouse SCC cells. (B) 10-point dose-response screen and IC50 determination for the KAT2B inhibitor GSK-4027 in SCC cells. (C) 10-point dose-response screen with the KAT2B/P300/CBP multi-target inhibitor Garcinol in SCC cells. (D) 10-point dose-response screen and IC50 determination for the CBP/P300 dual inhibitor SGC-CBP30. (E) 10-point dose-response screen with the KAT6A inhibitor WM-8014, examining its efficacy against KAT6A in SCC. (F) 10-point dose-response screen with another KAT6A inhibitor WM-119, providing a comparative analysis of KAT6A inhibition. (G) 10-point dose-response screen and IC50 determination with another PAK1 inhibitor NVS-PAK1, providing a comparative analysis of PAK1 inhibition. (H) Average IC50 values for selective histone acetyltransferase (HAT) inhibitors targeting the top significantly depleted genes identified in the RESTRICT-seq AZD4547 screen. This includes comparisons between SC-200891, GSK-4027, and SGC-CBP30, along with negative enantiomeric control compounds GSK4028, across human (A431, JHU-029, MR041718, SCC-25) and mouse (MR041718, 392-1). Bar graphs represent the mean ± SD. *n* = 3 technical replicates from three separate experiments. Statistical significance was determined using a two-sided Student’s unpaired t-test (*: p < 0.05, **: p < 0.01, ***: p < 0.001, ****: p < 0.0001).

**Supplemental Figure 12.**
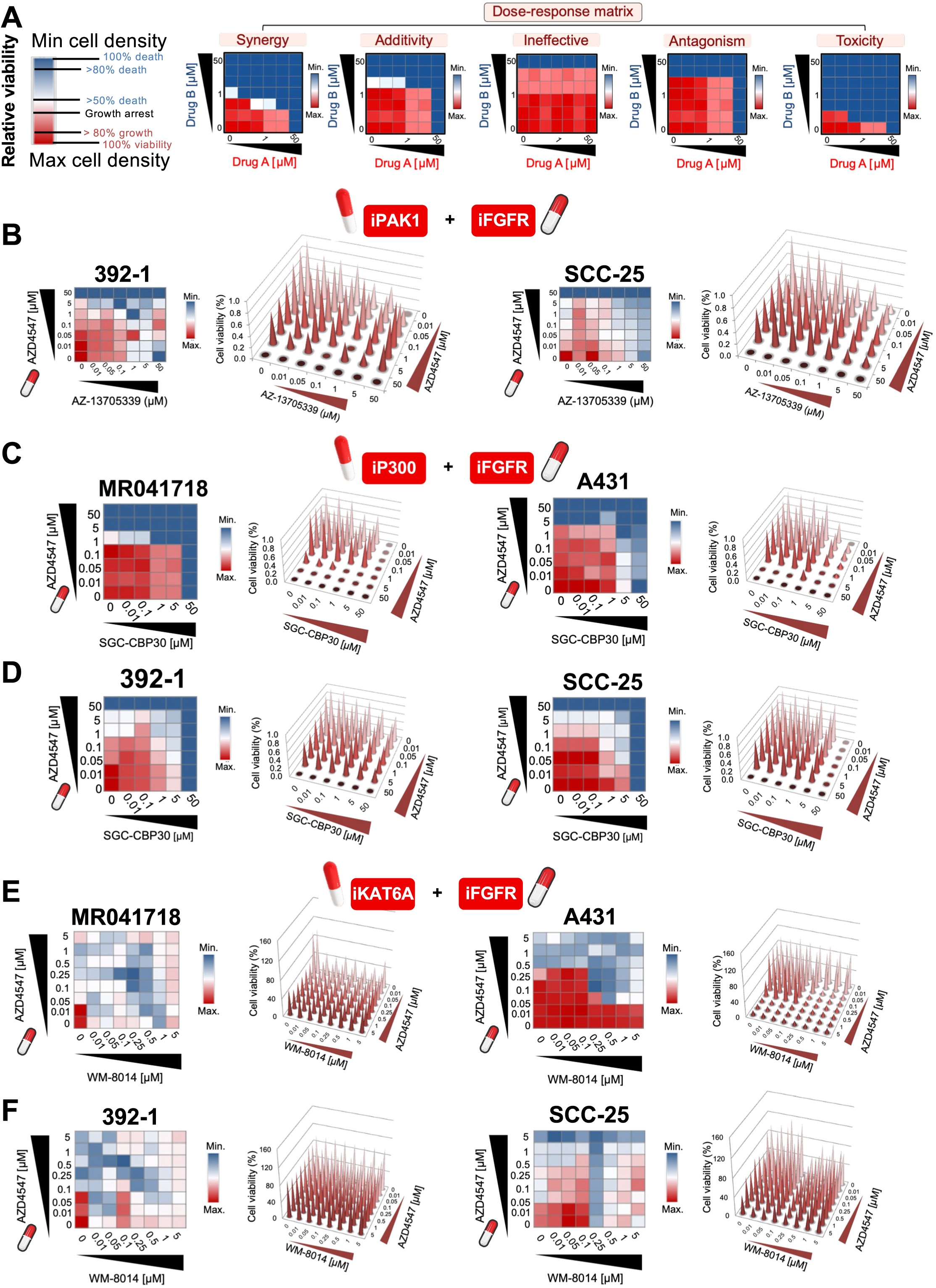
Synergistic inhibition of histone acetyltransferases results in predominantly additive reduction in tumor cell survival. (A) Schematic overview of synergistic dose-response small molecule inhibitor screens, illustrating the experimental setup for assessing combinatorial drug effects. (B) Synergistic lethality matrix depicting the effects of AZD4547 combined with PAK1 inhibitor AZ13705339 in mouse (392-1) and human (SCC-25) cSCC cells. (C-D) Synergistic lethality matrix illustrating the effects of P300 inhibitor SGC-CBP30 in combination with AZD4547 in mouse (392-1, MR041718) and human (A431, SCC-25) cSCC cells. (E-F) Synergistic lethality matrix showing the effects of KAT6A inhibitor WM-8014 with AZD4547 in mouse (392-1, MR041718) and human (A431, SCC-25) cSCC cells. *n* = 3 technical replicates from three separate experiments.

